# Lymph node contraction links sex-biased naive CD8 T cell decline to compromised antigen recognition during middle age

**DOI:** 10.1101/2025.02.11.637693

**Authors:** Lutz Menzel, Maria Zschummel, Meghan J. O’Melia, Mårten Sandstedt, Hengbo Zhou, Pin-Ji Lei, Lingshan Liu, Hang Lee, Debattama R. Sen, Lance L. Munn, Lena Jonasson, Timothy P. Padera

**Affiliations:** Department of Radiation Oncology, Edwin L. Steele Laboratories, Massachusetts General Hospital and Harvard Medical School, Boston, USA; Immunology of Aging, Leibniz Institute on Aging - Fritz Lipmann Institute e.V. (FLI), Jena, Germany; Harvard Medical School, Boston, USA; Broad Institute of Harvard University and Massachusetts Institute of Technology, Cambridge, USA; Krantz Family Center for Cancer Research, Massachusetts General Hospital, Harvard Medical School, Charlestown, MA, USA; Department of Health, Medicine and Caring Sciences, Linköping University, Linköping, Sweden; Clinical Department of Radiology, Region Östergötland, Linköping, Sweden; Center for Medical Image Science and Visualization (CMIV), Linköping University, Linköping, Sweden; Biostatistics Center, Harvard Medical School, Massachusetts General Hospital, Boston, Massachusetts; Clinical Department of Cardiology, Region Östergötland, Linköping, Sweden

## Abstract

The numerical abundance of diverse naive CD8 T cell clones is essential to provide broad protection against infection and cancer. Here, we uncover a sex-biased mechanism of immune aging in which an early, male-biased depletion of naive CD8 T cells driven by accelerated, antigen-agnostic differentiation into virtual memory cells is combined with age-related thymic involution that limits naive CD8 T cell replenishment. These mechanisms lead to contraction of lymph nodes and reduced local naive T cell clone availability, limiting cancer antigen recognition capacity. Therapeutic thymus regeneration via androgen ablation repopulates naive CD8 T cells in lymph nodes, which reinvigorates cancer-specific T cell responses and enhances responsiveness to immune checkpoint blockade. These findings reveal the crucial impact of sex and age on naive T cell clone abundance in lymph nodes and suggest strategies to restore immune competence in middle-aged males.

## Main

Age is a dominant risk factor for cancer, with individuals over 50 accounting for 90% of cancer-related mortality. Notably, males exhibit nearly twice the incidence and mortality risk compared to females^1,2^. While recent work has begun to clarify how cancer-targeting T cell responses are shaped by biological sex^3–5^ or age^6–8^, the intersection of these factors remains incompletely understood. Addressing these gaps may reveal strategies to restore immune competence and improve cancer outcomes across aging populations in a sex-specific manner.

The immune system’s ability to recognize and respond to a wide array of antigens is directly linked to the number of naive T cells and their clonotypic diversity. With age, the naive T cell compartment declines due to progressive thymic involution and ongoing peripheral attrition via differentiation into memory and senescence-prone effector cells (**Extended Data Fig. 1a**)^9–11^. The decline of naive T cells has been linked to reduced vaccine responses, impaired tumor surveillance, and reduced efficacy of immunotherapies^9,12,13^. Lymph nodes (LNs) are the primary sites for initiating immune responses to locoregional antigen sources, rendering them essential for effective anti-cancer immunity^14,15^. A decline in naive T cells reduces their local abundance within LNs, thereby limiting encounters between antigen-presenting cells (APCs) and cognate naive T cell clonotypes. These encounters are critical to generating adaptive immune responses and limiting them reduces antigen specific adaptive immunity^16^. Recent work has begun to clarify why the numerical availability of naive CD8 T cells in draining LNs is crucial for anti-cancer immunity. Anti-cancer immune responses are inherently stochastic and driven by rare, functionally critical T cells. As the size of the naive T cell precursor pool available for priming decreases, the probability of these rare cells being available for activation also decreases^15,17^. The primary source of naive T cells is the thymus, whose output is robust early in life through adolescence, establishing a diverse and self-tolerant peripheral T cell repertoire. While thymic involution has long been considered a feature of aging with limited impact on the adult immune system^18^, recent human data suggest it has active clinical relevance leading to higher long-term risk of mortality and incidence of cancer^19–21^. These data indicate that sustained thymic activity contributes to ongoing immune competence even later in life. How the decline in naive T cells and thymic atrophy throughout life influence the ability of LNs to initiate adaptive anti-cancer immune responses, and whether and how this process differs by sex-particularly during middle age (**Extended Data Fig. 1b**) when cancer incidence increases-remains unclear.

In this study, we bridge this gap by analyzing naive CD8 T cell dynamics across the lifespan in female and male mice, complemented by supporting observations from human datasets. Our data show a more rapid decline in naive CD8 T cell populations in males compared to females, leading to biological significantly fewer naive CD8 T cells in middle-aged males. Further, our data indicate that thymic involution alone is insufficient to explain the early and sex-biased loss of naive CD8 T cells; instead, peripheral antigen-agnostic differentiation into virtual memory cells acts as an amplifying attrition mechanism as thymic output becomes limiting. The differentiation into virtual memory T cells, which are shorter-lived and more prone to senescence^22^, reduces the abundance of unique naive T cell clones in LNs and is associated with impaired cancer recognition. Experimental thymus regeneration in male mice was able to restore the source of naive CD8 T cells and improved cancer-targeted responses in middle-aged, but not aged, mice. These mechanisms uncover the potential for immune regeneration strategies to mitigate age- and sex-related differential treatment responses to immune checkpoint blockade. Understanding this mechanism is pivotal for developing novel therapeutic approaches to mitigate immune aging, particularly by addressing sex-specific vulnerabilities in thymus and naive T cell regeneration^23–25^.

## Results

### Age and sex are critical factors for tumor draining LN-derived T cell responses to melanoma

Analysis of data from the Surveillance, Epidemiology, and End Results (SEER)^26^ program indicates that males have higher overall cancer incidence rates, including melanoma, with these differences becoming apparent in middle age **(Fig. 1a, Extended Data Fig. 1b)**. Both sexes had similar involvement of tumor-infiltrating T cells (TILs) in melanoma across age groups based on transcriptional immune scores **(Extended Data Fig. 1c)**. However, defining T cell differentiation states using transcriptional module scoring of canonical gene signatures **(Extended Data Fig. 1d-e)** in published samples from melanoma patients^27–30^ revealed a higher relative proportion of naive-like CD8 T cells to antigen experienced (cytotoxic and exhausted) T cells within tumors from male patients compared to females. These data suggest restricted antigen specific activation in males **(Fig. 1b,c)**, which could be a result of deficits in T cell priming in tumor-draining LNs (tdLNs). In addition to well-documented sex and age-related T cell impairments, such as exhaustion, dysfunction or the development of suppressor phenotypes^3–5,7,8^, a notable proportion of TILs in older individuals retain a naive-like state. Consequently, these TILs may not actively engage in, and could remain bystanders to, antitumor immune responses^29,31–34^.

**Fig. 1.**
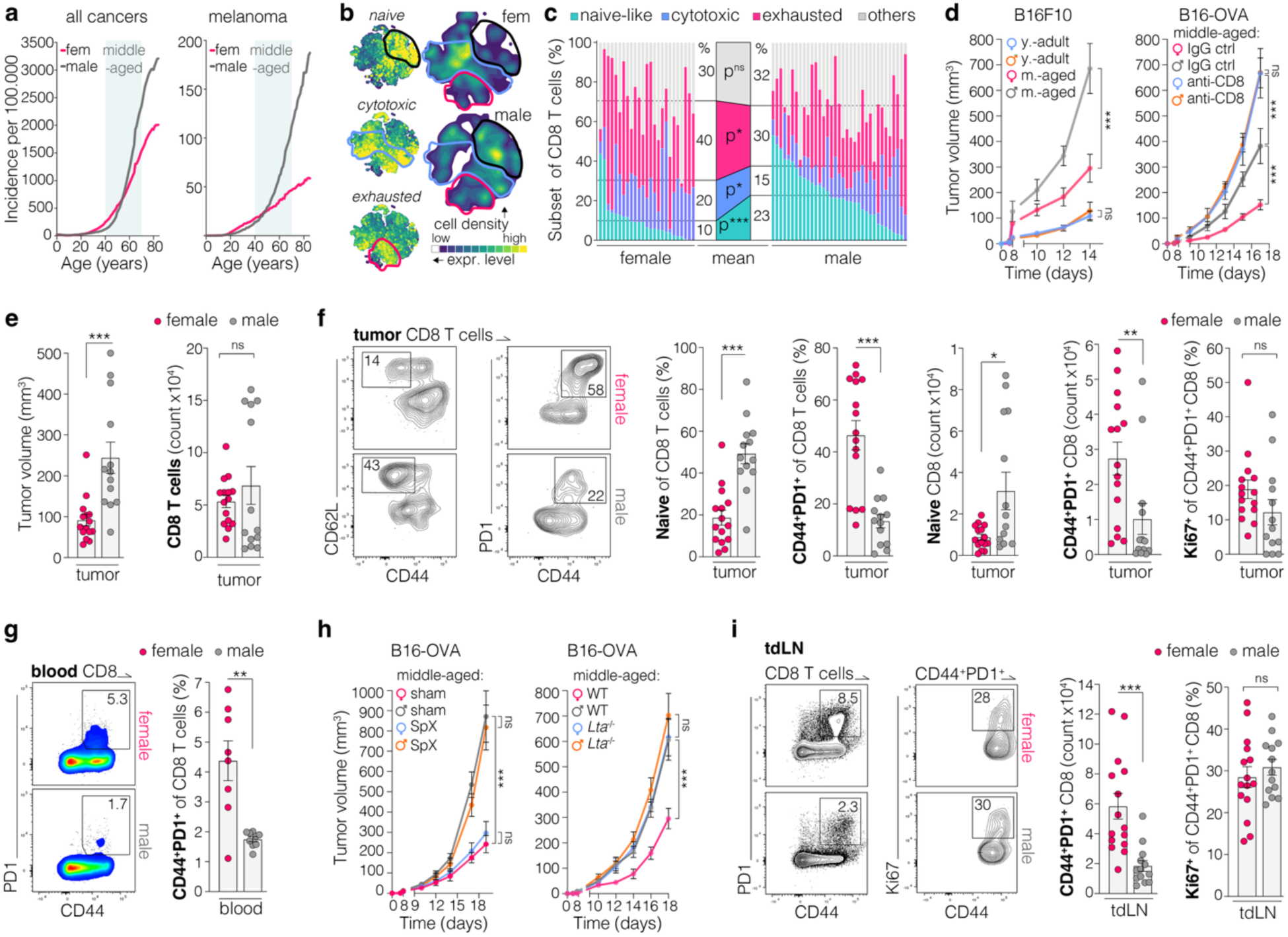
Age and sex are critical factors for tdLN-derived T cell responses to melanoma. **a,** incidence per 100,000 individuals, data from SEER^26^ (n=1.8-17.8x10^6^ cases per age and sex group). **b,** cell density projections and expression levels of gene module scores of CD8 T tumor infiltrating lymphocytes (TILs) isolated from melanomas, integrated public data sets^27–30^. Corresponding data in **Extended Data Fig. 1b-e**. **c,** frequency of CD8 TIL subsets in individual patients and in average per sex (females n=25, males n=35). **d,** B16F10 (left) tumor growth in untreated young adult (y.-adults, n=5) or middle-aged (m.-aged, n=8) mice . B16-OVA (right) tumor growth in middle-aged mice with and without CD8 T cell depletion (n=5/group). **e,** B16-OVA tumor volume and CD8 T cell infiltration 13-15 days after implantation (corresponding to **f,g**)**. f,** flow cytometry analysis of CD8 T cells isolated from B16-OVA tumors. Gating strategy in **Extended Data Fig. 3a**. **g,** CD8 T cells from peripheral blood of middle-aged mice with B16-OVA tumors. **h,** B16-OVA tumor growth in middle-aged mice, *left*, after sham surgery or splenectomy (SpX), n=8/group, and *right*, in wild-type (WT) or lymphotoxin alpha knock out (*Lta*^-/-^) mice, n=8-10/group. **i,** Flow cytometry analysis of CD8 T cells from tumor-draining lymph nodes (tdLNs) of middle-aged mice with B16-OVA tumors 13-15 days after implantation. Gating strategy in **Extended Data Fig. 3a**. Data shown as mean ± SEM, each data point representing a single LN or tumor from an individual mouse or person. Statistics calculated with Mann-Whitney U test (MWU) test (c,e,f,g,i) or two-way repeated measures ANOVA (d,h) to assess overall effects across time and group; ns=nonsignificant, *p<0.05, **p<0.01, ***p<0.001.

To assess CD8 T cell-mediated tumor control across age and sex, we compared tumor growth in young adult (7-10 weeks) and middle-aged (9-12 month) mice (**Extended Data Fig. 1b)** using the B16F10 melanoma model. In young adults, males and females showed comparable tumor growth. By middle age, however, males displayed a marked loss of tumor control, whereas females retained substantially lower tumor growth. To directly test CD8 T cell dependence, we performed CD8 T cell depletion experiments in the B16-OVA melanoma model, which enables tracking of antigen-specific CD8 T cell responses. CD8 depletion accelerated tumor growth in both sexes but abolished all sex-related differences, demonstrating that the divergence in tumor control is CD8 T cell dependent **(Fig. 1d, f Extended Data Fig. 2b-d)**. Consistent with this, middle-aged females mounted a more robust response to anti–PD1 therapy than middle-aged males, further supporting a sex-biased T cell contribution to tumor restraint **(Extended Data Fig. 2e)**. At an early stage of tumor growth, when significant sex-related differences in tumor size were already apparent **(Fig. 1e *left*)**, we performed flow cytometric analysis of TILs in B16-OVA tumors from middle-aged mice. Consistent with patient data **(Fig. 1b,c),** tumors in middle-aged females contained a higher number of antigen-experienced (CD44^+^PD1^+^) and tetramer positive T cells and T cells with functional effector phenotype (Granzyme-B^+^), while males exhibited a predominance of naive CD8 T cells **(Fig. 1f; Extended Data Fig. 3b)**. This disparity occurred despite comparable overall CD8 T cell infiltration and intratumoral proliferation of effector T cells **(Fig. 1e *right*, f)**. The number of exhausted T cells (TIM3^+^PD1^+^) was higher in females and regulatory T cells similar between sexes **(Extended Data Fig. 3c)**. The relative increase in naive T cells in tumors from middle-aged males suggests that they are not properly primed prior to entering the tumor. The ability of naive T cells to enter into the primary tumor is likely a result of abnormal tumor vasculature, including dysregulation of adhesion molecules on the endothelial cells leading to aberrant lymphocyte trafficking^35^.

The higher frequency of CD44⁺PD1⁺ CD8 T cells was also present in the blood of middle-aged female mice, suggesting that these cells are activated outside the tumor and likely reflect more effective priming in LNs compared to males **(Fig. 1g)**. We performed splenectomy (SpX) or utilized lymphotoxin alpha-deficient (*Lta*^-/-^) middle-aged mice—which lack functional LNs^36^—and found that LNs, rather than the spleen, are critical for supporting the superior tumor recognition capacity in middle-aged females **(Fig. 1h)**. Consistently, tdLNs from middle-aged female mice contained significantly higher numbers of antigen-experienced (CD44^+^PD1^+^) and tumor specific (tetramer^+^) CD8 T cells, independent of the proliferation capacity of activated cells **(Fig. 1i, Extended Data Fig. 3d,e)**, suggesting a higher probability of naive T cells that recognize tumor antigens in tdLNs of middle-aged females. Together, these findings indicate that tumors in middle-aged females have a higher proportion of cancer-targeting CD8 effector T cells that are primed in tdLNs before they infiltrate the tumor, consistent with the human data (**Fig. 1b-c**).

Of note, while sex disparities of intradermal APCs have been reported for adult mice^37^, LNs in middle-aged humans did not contain dysregulated proportions of APCs **(Extended Data Fig. 4a)**. Further no differences in antigen uptake or presentation between the sexes in middle-aged mice was observed, as assessed using the B16-SIINFEKL-zsGreen tumor model and DQ-OVA immunization **(Extended Data Fig. 4b-e)**. This lack of sex-based differences in antigen presentation in LNs prompted us to test whether middle-aged males possess a restricted naive T cell pool and, consequently, contained limited numbers of unique TCRs capable of recognizing tumor antigens in LNs.

### Age-related decline of naive CD8 T cells and contraction of LNs occur earlier in males

Volume segmentation of full-body computed tomography (CT) scans of humans from a publicly available database^38,39^ revealed larger spleens but significantly smaller LNs in males compared to females **(Fig. 2a; Extended Data Fig. 5a,b)**. With age, LNs can accumulate fat due to lipomatosis, disrupting the correlation between volume and cellularity, which can potentially lead to an underestimation in the decline in cellularity^40^. In middle-aged mice, LNs did not occur with a dysregulated stromal structure **(Extended Data Fig. 6a)**. Consistent with human LN data, murine LN size and cellularity declined with age, occurring earlier and more markedly in males, and resulting in a significant sex disparity by middle age **(Fig. 2b,c)**. The differences were more pronounced in skin-draining compared to mesenteric LNs and were independent of reproductive status and genetic background **(Extended Data Fig. 6b-d)**. LN weight strongly correlated with cellularity during physiological aging and under lymphopenic conditions, emphasizing the co-dependence of LN size with the number of lymphocytes **(Fig. 2d; Extended Data Fig. 6e)**.

**Fig. 2.**
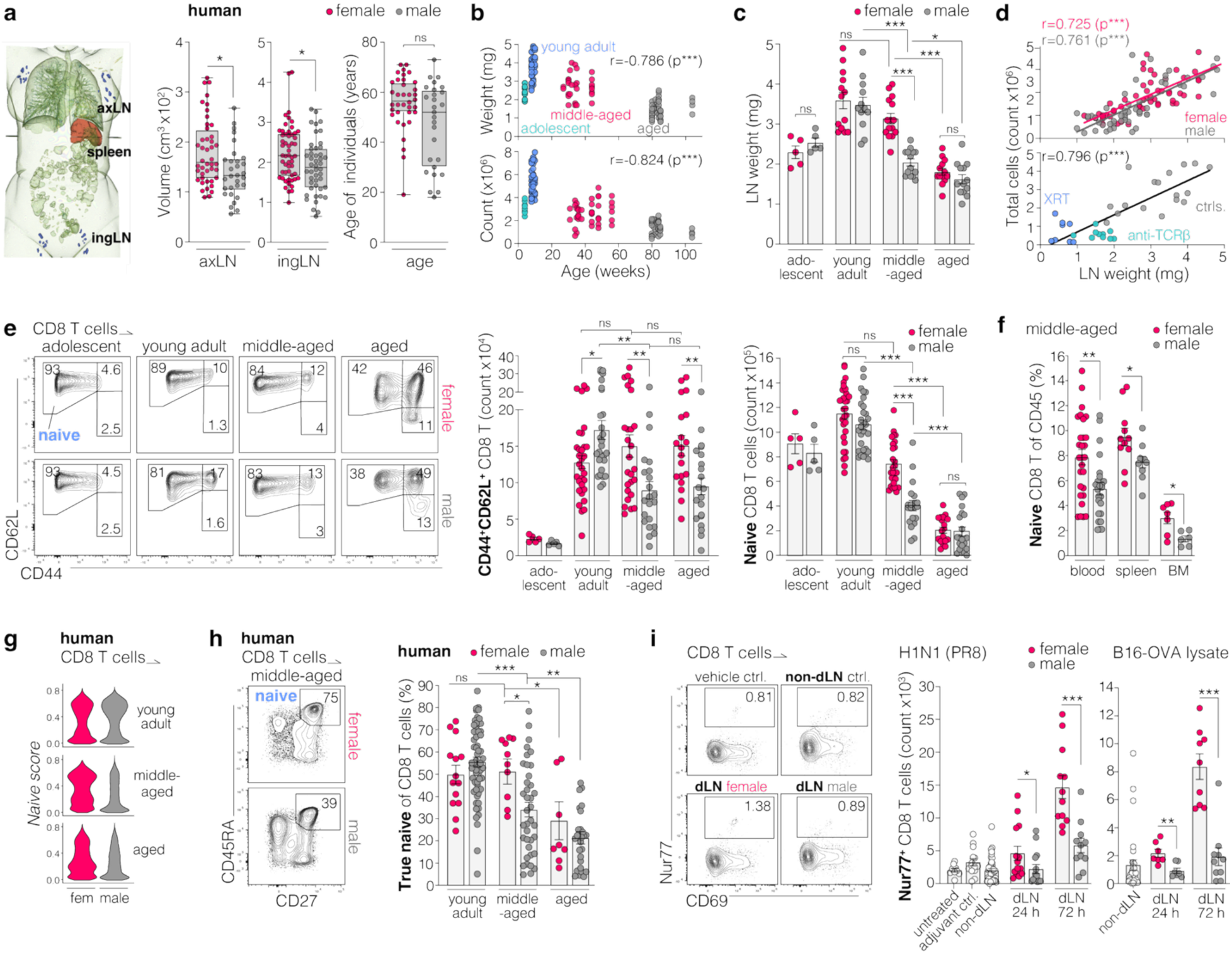
Age-related naive CD8 T cell decline and LN contraction occurs earlier in males. **a,** Average axillary (axLN) and inguinal (ingLN) LN volume per human individual, segmented from computer tomography (CT) scans^38,39^, age of individuals (when available, for others, age is unspecified) in corresponding data in **Extended Data Fig. 5a,b**. **b,** Correlation of individual ingLN weight (top) or cellularity (bottom) with age in female and male mice. Adolescent mice are shown but excluded from correlation calculations. **c,** LN weights in age-groups of female/male mice. **d, top**, Correlation of LN weight with LN cellularity across all ages. **Bottom**, Correlation after irradiation (XRT, 4 Gy) or after T cell depletion with anti-TCRβ antibody. **e,** Flow cytometry analysis of naive (CD44^-^CD62L^+^) and memory (CD44^+^CD62L^+^) CD8 T cells from ingLNs of mice. Gating strategy in **Extended Data Fig. 6f**. **f,** Frequency of naive CD8 T cells in the blood, spleen and bone marrow (BM) of middle-aged mice. **g,** Expression levels of naive T cell genes (*Naive score*) in human CD8 T cells, ABF300 study^46^. Naive score described in **Extended Data Fig. 5c**. **h,** flow cytometry quantification of true naive CD8 T cells from human PBMCs, public data set^47^. Gating Strategy in **Extended Data Fig. 5g**. **i,** Nur77 expression in CD8 T cells from the draining lymph node (dLN) of middle-aged mice 24 or 72 h after immunization with UV light inactivated H1N1 (PR8) influenza virus or B16F10-OVA melanoma cell lysate. Data shown as mean ± SEM with data points representing a single LN from an individual mouse or person. Statistics calculated with MWU test (a,c,e,f,h,i) or Spearman correlation (b,d) and trend lines are shown for visualization (d), ns=nonsignificant, *p<0.05, **p<0.01, ***p<0.001.

Previous studies have established that the naive CD8 T cell compartment undergoes substantial remodeling with age, including numerical reductions in naive CD8 T cells^41–44^. Much of this decline has traditionally been viewed in the context of a proportional shift from naive to memory phenotypes^9,45^. However, in our data we observe quantitative stability or loss of CD44⁺CD62L⁺ memory-phenotype cells and a clear loss of naive CD8 T cells in inguinal LNs (ingLN) beginning shortly after young adulthood, with naive cells showing a particularly pronounced reduction in males compared to females, declining from ∼10^6^ cells/ingLN in young adults of both sexes to ∼8x10^5^ cells/ingLN in middle-aged females and ∼4x10^5^ cells/ingLN in middle-aged males, with both sexes converging with advanced age **(Fig. 2e)**. This decline is not restricted to LNs but is also evident in naive CD8 T cell frequency in blood, spleen, and bone marrow (BM) **(Fig. 2f)**, underscoring a systemic loss of naive CD8 T cell clones with advancing age. Published multimodal datasets of human T cells from blood and LNs^46–48^ corroborated these findings, showing a decline of naive CD8 T cells determined either by flow cytometry or by applying a gene expression “*Naive score*” on scRNA-seq data **(Fig. 2g,h; Extended Data Fig. 5d,e)**.

To examine TCR diversity in LNs, we performed TCR sequencing of CD8 T cells from LNs of middle-aged mice and found a maximally diverse TCR repertoire with over 96% of unique T cell clonotypes in non-reactive LNs independent of sex, mirroring observations in humans^48^ **(Extended Data Fig. 7a-e)**. Thus, we hypothesized that the absolute number of naive T cells within a LN at any given time is directly correlated with the number of unique TCRs present and is therefore a rate-limiting factor for the probability of antigen recognition. To test this, we immunized middle-aged mice intradermally in the flank with either inactivated H1N1 virus particles or B16-OVA tumor cell lysates and analyzed the draining LNs at 24- and 72-hours post-immunization to assess early T cell activation. Both sexes exhibited similar frequencies of CD8 T cells expressing the general activation marker CD69. However, a significantly greater number of CD8 T cells in females exhibited TCR-specific activation, as indicated by a higher proportion of Nur77⁺ cells^49^ **(Fig. 2i; Extended Data Fig. 7f)**. We attribute this sex imbalance to the partial depletion of naive CD8 T cells in middle-aged males and the concomitant contraction of LNs, which are central to locoregional antigen processing and recognition. These data highlight the functional impact of higher naive T cell numbers in middle-aged females on the generation of antigen specific immune responses. These findings support a model in which reduced availability of naive CD8 T cells limits the probability of recruiting cancer-reactive clones. This is consistent with recent work^15,17^ showing that a diminished naive T cell pool is expected to directly constrain the formation and diversity of these precursor populations, thereby reducing the likelihood of effective cancer-specific T cell responses. The greater activation of CD8 T cells in LNs of middle-aged female mice is consistent with an increase in the proportion of antigen-experienced CD8 TILs in the human data **(Fig. 1d-e)**. Even though middle-aged male mice have fewer total numbers of naive CD8 T cells, they also have fewer antigen experienced CD8 T cells due to a reduced numerical abundance of TCR clones in LNs. Thus, as a proportion, the TILs in the human data remain skewed toward a naive-like phenotype in males.

### Enhanced propensity for an antigen-agnostic loss of T cell naivety in males

While thymic involution is a well-known feature of aging, its functional relevance remains debated^18,20^, as peripheral T cell numbers, particularly in the CD4 compartment, remain relatively stable in adulthood (**Extended Data Fig. 5f)**, consistent with previous reports^50^. However, other studies^41–44^ and our data reveal a progressive, selective decline in naive CD8 T cells with age, suggesting that peripheral maintenance is insufficient to compensate once thymic output declines with age. This raises the question of what drives this loss and why it occurs earlier in males.

Naive CD8 T cells are long-lived but can exit the naive pool through activation into effector and central memory (Tcm) cells upon antigen encounter, or via antigen-agnostic differentiation into virtual memory (Tvm) cells (**Fig. 3a**)^51–54^. While the effector/Tcm route can be pathogen-driven^9,13,55^, our data show that naive CD8 T cells still decline with age in mice housed in low-pathogen conditions **(Fig. 2e,f)**. CD44⁺CD62L⁻ effector-like cells did not differ by sex across the lifespan, suggesting similar antigen exposure. However, young adult males exhibit an increased frequency of CD44⁺CD62L⁺ CD8 T cells already before any sex difference in LN cellularity is apparent in middle-aged (**Extended Data Fig. 8a**). In line with previous studies^22,56,57^, most CD44⁺CD62L⁺ cells were not true Tcm (CD44⁺CD49d⁺), but Tvm cells (CD44⁺CD122⁺CD49d⁻) **(Fig. 3b; Extended Data Fig. 8b)**. The propensity of naive CD8 T cells to undergo antigen-agnostic differentiation into Tvm cells is shaped by their clone-specific sensitivity to self-peptide-MHC and is reflected by CD5 expression, which is stably imprinted during thymic selection^53,58,59^. CD5^high^ cells are more self-reactive and more likely to undergo Tvm conversion in the periphery over time^51,53^.

**Fig. 3.**
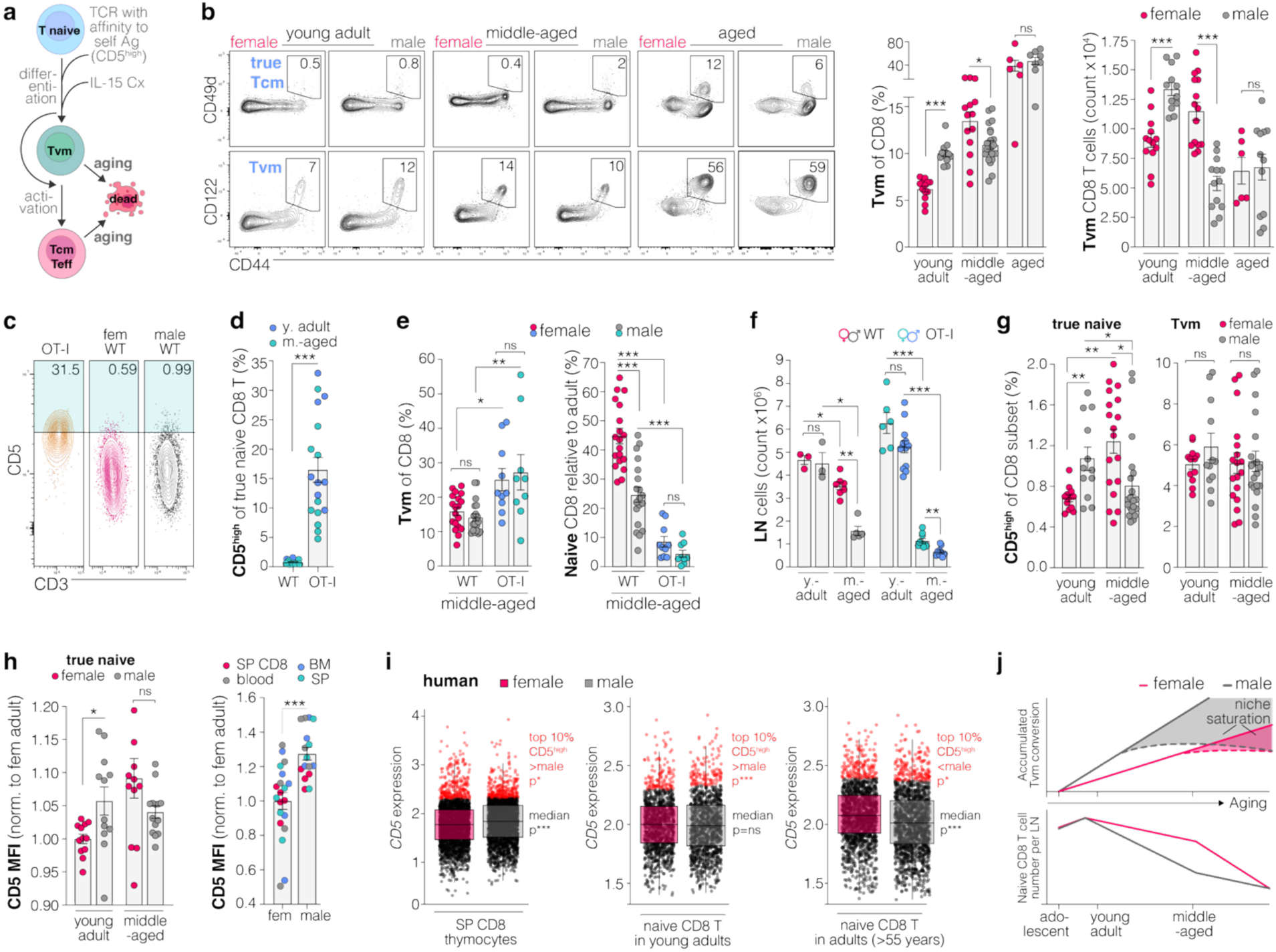
Enhanced propensity for antigen-inexperienced loss of T cell naivety in male mice. **a,** schematic illustration of naive T cell differentiation into virtual memory (Tvm), effector (Teff) and central memory (Tcm) cells; T cell receptor (TCR), IL-15/IL15 receptor alpha complex (IL-15 Cx). **b,** Central memory T cells (Tcm) and Tvm cells in ingLNs of WT mice. Gating strategy in **Extended Data Fig. 6f**. **c,** Example plots and, **d**, quantification of CD5^high^ cells in true naive CD8 T cells from ingLNs of WT or OT-I mice. OT-I mice were used as a known positive control. **e,** Quantification of Tvm and naive CD8 T cells in middle-aged WT and OT-I mice. **f**, Quantification of total cell counts of ingLNs of young adults (y. adult) and middle-aged (m.-aged) WT and OT-I mice. **g,** Quantification of CD5^high^ cells from ingLNs of WT as a percent of true naive CD8 T cells or Tvms. **h,** CD5 mean fluorescence intensity (MFI) normalized to young adult female mice and MFI of CD5 single positive CD8 thymocytes and true naive CD8 T cells from blood, bone marrow (BM) and spleen (SP). **i,**Human CD5 expression in top 10% CD5^high^ cells and median overall CD5 expression in, *left*: CD8 SP thymocytes from young adults or, *right*: naive CD8 T cells in young adults or adults over 55 years of age. Data from published datasets of human thymocytes^60,61^ and PBMCs^46^. **j,** Illustrative projection of cumulative conversion of CD5^high^ naive CD8 T cells into Tvm cells during aging. ***Top***, indicated are the linear accumulation of Tvm cells over time (solid line) and the assumed deceleration in accumulation due to competition and the prior conversion of naive CD8 T cells with highest CD5 levels (dashed line: slow down). ***Bottom***, the Tvm formation associated attrition of naive CD8 T cells. Data shown as mean ± SEM, each data point representing a single LN or tumor from an individual animal. Statistics calculated with MWU test (b,d,e,f,g,h), Wilcoxon rank-sum test (i), ns=nonsignificant, *p<0.05, **p<0.01, ***p<0.001.

To test whether high self-reactivity alone is sufficient to drive this process, we used the OT-I mouse model, in which all CD8 T cells express monoclonal TCRs and have elevated reactivity for endogenous self-antigens^51,53^, resulting in higher proportions of CD5^high^ cells compared to WT mice **(Fig. 3c,d, Extended Data Fig. 8c)**. As expected, OT-I mice exhibited accelerated Tvm formation, a pronounced reduction of naive CD8 T cells in LNs with age, and reduced cellularity of both LNs and spleen in middle-age of both sexes **(Fig. 3e,f; Extended Data Fig. 8d)**. These findings confirm that high TCR self-reactivity is sufficient to drive antigen-agnostic Tvm differentiation and naive CD8 T cell pool contraction. To determine whether this mechanism operates in a physiologically diverse TCR setting, we analyzed WT mice and found that CD5^high^ cells were enriched in the Tvm population relative to naive CD8 T cells. CD5^high^ true naive CD8 T cells were more frequent in young adult males than females **(Fig. 3g)**, accompanied by higher CD5 expression levels across LNs and other organs **(Fig. 3h)** suggesting a sex-biased predisposition to early Tvm conversion. In mice, this difference reversed with age as CD5^high^ naive CD8 T cell frequencies increased in females but declined in middle-aged males, likely reflecting prior selective loss of self-reactive clones through Tvm differentiation in middle-aged males **(Fig. 3g)**. Consistent with the increased frequency of CD5^high^ cells in young male mice, our analysis of publicly available scRNA-sequencing data^47,64,65^ from human SP CD8 thymocytes and naive CD8 T cells in PBMCs showed elevated CD5 expression among the top 10% CD5^high^ cells in males compared to females **(Fig. 3i)**.

Although the difference in CD5^high^ naive CD8 T cell frequency between young adult females and males is modest (approximately 0.7% vs. 1.1%), these cells arise from a compartment comprising >85% of all CD8 T cells, resulting in a substantial absolute pool of higher self-reactive clones available for Tvm conversion. To assess the long-term impact, we implemented a simple model of naive-to-Tvm conversion with sex-specific rates based on the CD5^high^ naive CD8 T cell frequency **(Supplementary Fig. 1a-d)**, showing that small initial differences in CD5^high^ frequency can drive cumulative divergence in Tvm generation and naive CD8 T cell depletion over time as shown in in the conceptual trajectories shown in **Fig. 3j**.

### Transient experimental Tvm induction initiates a naive CD8 T cell decline in long-term

From an evolutionary perspective, the early skimming of naive CD8 T cells to Tvm may reflect a functional trade-off. Tvm cells offer rapid, cytokine-driven responses that prioritize short-term immune readiness over long-term repertoire maintenance^51,63–65^. Compared to naive cells, Tvm cells display enhanced effector cytokine production upon stimulation, yet lose proliferative capacity, upregulate senescence-associated transcriptional signatures **(Fig. 4a-c, Extended Data Fig. 9a-c)**, and increase apoptosis rates with aging (**Extended Data Fig. 9d)** with minimal age and sex dependent metabolic differences **(Supplementary Fig. 2)**. LNs continue to shrink with age despite Tvm formation, as Tvm cells have a shorter lifespan due to increased apoptosis^22,62^, and preferentially relocate to the spleen, bone marrow **(Extended Data Fig. 9e)** or liver^51^. Both of these processes contribute to a sex-biased loss of naive CD8 T cells and LN contraction earlier in life.

**Fig. 4.**
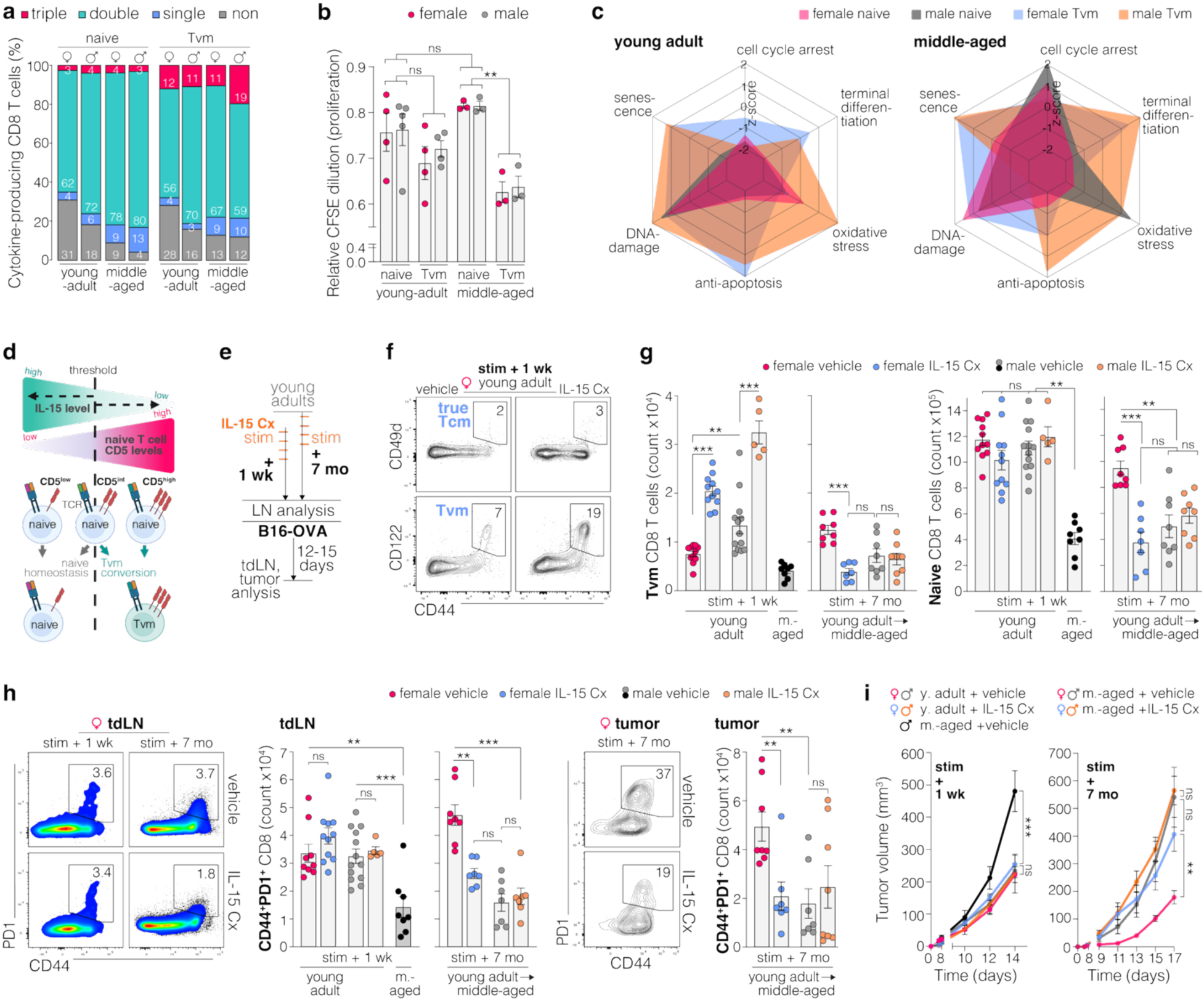
Experimental induction of Tvm in young females leads to restricted naive CD8 T cell pool in middle-aged. **a,** Polyfunctional cytokine production in subsets of CD8 T cells after in vitro stimulation, showing the proportion of cells producing combinations of IL-2, IFNγ, and TNF as measure for effector capacity. **b**, Proliferation capacity of CD8 T cell subsets after in vitro stimulation, quantification depicted by dilution of CFSE MFI relative to unstimulated control. **c**, Radar plots showing z-scored mean expression of senescence-associated gene sets in young adult and middle-aged naive and Tvm CD8 T cells across all groups. Gene sets described in material and methods. **d**, Schematic model of naive-to-Tvm conversion. IL-15 acts as a permissive signal that establishes a self-reactivity threshold (CD5 level, here depicted as either low, intermediate (int) and high). Only naive CD8 T cells above this threshold undergo conversion to Tvm. **e,** Experimental setup, young adult and middle-aged mice treated with vehicle or IL-15/IL-15 receptor alpha complex (Il-15 Cx once weekly for 4 times) with LN analysis and B16-OVA tumor implantation 1 week after the acute stimulation phase (stim + 1 wk) or 7 months later (stim + 7 mo). **f,** Representative flow cytometry plots. **g,** Quantification of CD8 T cells in ingLNs of mice treated with vehicle or IL-15 Cx as described in **f**; middle-aged (m.-aged). **h,** Flow cytometry analysis of CD8 T cells from the tdLNs or the primary tumor, young-adults (y.-adult) and middle-aged (m.-aged). **i,** B16-OVA tumor growth in young adult and middle-aged mice treated with vehicle or IL-15 Cx as described in **a**, n=7-8/group. Data shown as mean ± SEM, each data point representing a single LN or tumor from an individual animal. Statistics calculated with Student’s t-test (b), MWU test (g,h), two-way repeated measures ANOVA (j), ns=nonsignificant, *p<0.05, **p<0.01, ***p<0.001.

To further establish that Tvm formation can lead to depletion of naive CD8 T cell populations, we used IL-15/IL-15 receptor alpha complexes (IL-15 Cx), the biologically most active form of IL-15, to experimentally drive Tvm formation in young adults. As illustrated in **Fig. 4d**, IL-15 does not directly induce Tvm differentiation but acts as a permissive co-stimulatory signal that lowers the activation threshold of self-reactivity (CD5^level^) of naive CD8 T cells for converting into Tvm cells^51,66–68^. Thus, IL-15 Cx enables experimental lowering of the self-affinity threshold for Tvm formation in a setting with preserved TCR diversity but low baseline conversion.

Recombinant IL-15 Cx was administered once weekly for 4 times to young adult mice **(Fig. 4e).** IL-15 Cx-treated females allowed us to ask whether transient, experimentally induced Tvm formation could mimic male-like immune aging. This approach allowed us to probe the causal link between premature Tvm skewing and later-life naive T cell loss in a setting where these features are not inherently coupled, thereby isolating the hypothesized process from confounding differences. We observed a marked expansion of Tvm cells in LNs of young adult female mice after the acute IL-15 Cx stimulation phase (stim + 3 wk). Remarkably, this transient intervention in female mice led to a significant and lasting reduction in naive CD8 T cells seven months later (stim + 7 mo) mimicking the profile observed in age-matched untreated male mice **(Fig. 4f,g)**. As Tvm cells are more prone to apoptosis and preferentially leave LNs for other organs, their accelerated conversion from naive CD8 T cells provides an explanation for the sustained depletion of CD8 T cell populations observed after IL-15Cx treatment. We next assessed whether endogenous IL-15 levels differ between sexes. IL-15 transcript levels were comparable at all ages, while protein levels showed modest decrease only in young males **(Extended Data Fig. 9f,g)**. As the transcription of IL-15 is not different in young males and females, the reduction in IL-15 in young males could be a result of increased IL-15 consumption by the expanding Tvm compartment. Further, these data indicate that sex-specific differences in IL-15 availability are unlikely to underlie the observed phenotype.

Although IL-15 Cx primarily expands pre-existing Tvm cells, it can also induce naive-to-Tvm conversion ^68^. Our data show that this transient expansion of the Tvm pool was sufficient to drive a long-lasting reduction in naive CD8 T cell numbers.

While the acute expansion of Tvm cells had no measurable effect on tumor rejection shortly after treatment (stim + 3 wk), early-life IL-15 Cx treatment led to decreased numbers of activated CD8 T cells within tdLNs and the tumor, and reduced tumor control in middle-aged female mice **(Fig. 4h,i)**. These findings demonstrate that transient induction of Tvm cells in young adult female mice preserves short-term anti-tumor effector function but accelerates long-term naive CD8 T cell attrition, thereby mimicking key features of male-biased immune aging.

### Early thymic involution in males limits naive CD8 T cell replenishment and facilitates LN contraction

Given the significant loss of naive CD8 T cells via Tcm and Tvm differentiation, we next asked why this decline is not adequately replenished over time. While no direct causal link has been established between thymic involution and Tvm differentiation, the increased abundance of Tvms in lymphopenic conditions and during aging likely reflects how progressive naive T cell loss makes thymic insufficiency functionally relevant^52,69,70^. Consistent with published data, we observed thymus involution in humans with significant sex differences, particularly in young adults and middle-aged individuals **(Fig. 5a)**, matching the decline in LN size and naive CD8 T cell numbers measured in both mice and humans **(Fig. 2e,h)**. Using data from a subsample of the SCAPIS cohort^71^, which included thymic health and naive CD8 T cell counts in middle-aged individuals, we showed a significant correlation between poor thymic health and lower numbers of naive CD8 T cells (**Fig. 5b**). Using data from Drabkin *et al.*^72^ and the SCAPIS cohort,^73^ we also showed age- and sex-related thymus involution using similar CT-based analysis methods **(Extended Data Fig. 10a-c)**. Across individuals, the thymus involution score and recent thymic emigrant (RTE) prevalence strongly correlated with the frequency of naive CD8 T cells, linking thymic atrophy to peripheral naive T cell supply **(Fig. 5b,c; Extended Data Fig. 10d)**. The lack of recovery from the sex-biased loss in naive CD8 T cells results from the poorer thymic health in males compared to age-matched females. Interestingly, while CD4 RTEs also declined with age and showed a similar sex bias, naive CD4 T cell numbers remained relatively stable **(Extended Data Fig. 10e)**, consistent with their lower conversion into memory-like cells and increased longevity^50^. In humans, naive CD4 T cell numbers remain stable through midlife, likely due to a compensatory reduction in CD4 T cell loss rate as thymic output wanes^50^, whereas naive CD8 T cells exhibit a more pronounced numerical decline due to Tvm conversion with thymic compensation insufficient to maintain their population^46,74^.

**Fig. 5.**
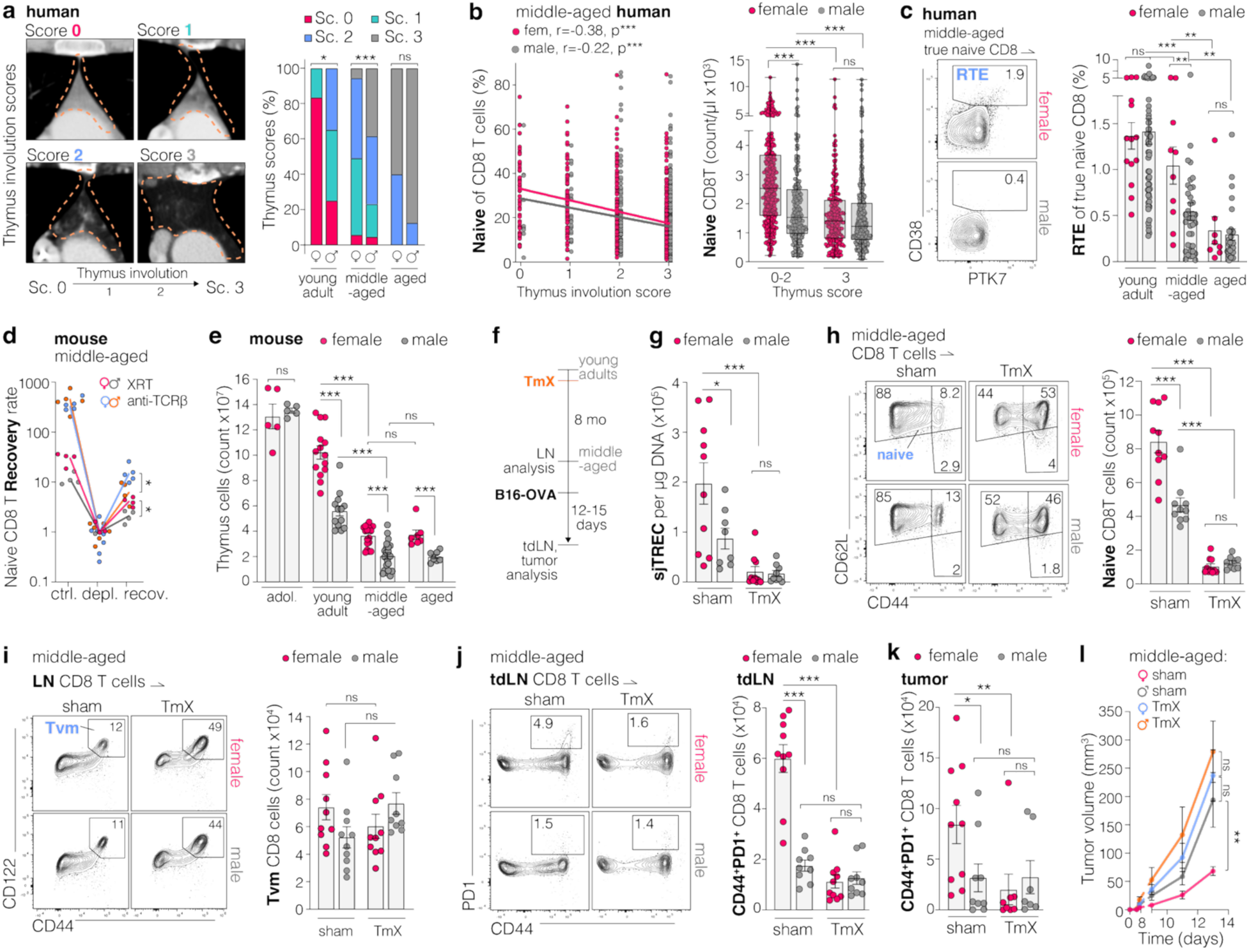
Early thymic involution in males limits naive CD8 T cell replenishment and facilitates LN contraction. **a,** Analysis of human thymus involution in CT scans, score (Sc.) 0: predominantly soft tissue; score 1: parity of fatty and soft tissue; score 2: predominantly fat; score 3: complete fatty replacement, Scoring system details in **Extended Data Fig. 10>a**. **b, *left*,** Correlation between the thymus involution score and naive CD8 T cell frequency and ***right***, number of naive CD8 T cells per µl of blood in individuals with preserved thymus (score 0-2) or fully involuted thymus (score 3). **c,** flow cytometry data from human PBMCs for recent thymic emigrants (RTE, CD38^high^) in true naive CD8 T cells (data from^47^). Gating strategy in **Extended Data Fig. 5f**. **d,** Recovery rate of naive CD8 T cells 7 days after irradiation (XRT) or anti-TCRß T cell depletion (depl.) and after a period of recovery (recov.): 28 days for XRT and 19 days for anti-TCRß. Cell numbers normalized to the mean at the depletion timepoint. e, Total thymocyte counts in male and female mice across the lifespan. **f,** Experimental setup, thymectomy (TmX) was performed in 5 week old mice and aged for 8 months until middle-aged, then a LN was analyzed and B16-OVA tumors were implanted. **g,** Signal joint T cell receptor excision circle (sjTREC) quantification in lymphocytes of LNs from middle-aged mice after sham surgery or TmX performed when young. **h,** Flow cytometry of CD8 T cells and quantification of true naive CD8 T cells and **i,** Tvm in middle-aged mice after sham surgery or TmX. Gating strategy in **Extended Data Fig. 6f**. **j and k**. Flow cytometry analysis of CD8 T cells from the tdlN **(j)** or primary tumor **(k). l,** B16-OVA tumor growth in middle-aged mice after sham surgery or TmX performed when 5 weeks old, n=9-10 mice/group. Data shown as mean ± SEM, each data point representing a single LN or tumor from an individual animal. Statistics calculated with MWU test (b *right*,c,d,e,g,h,i,j,k), Fisher’s exact test (a), Spearman correlation, trend lines are shown for visualization (b *left*), two-way repeated measures ANOVA (l); ns=nonsignificant, *p<0.05, **p<0.01, ***p<0.001.

Mechanistically, mouse studies recapitulated these observations. Mimicking the aging effect on LNs with experimental lymphodepletion, naive CD8 T cells recovered faster in middle-aged females than males **(Fig. 5d)**. The lymphopenia induced a temporary increase in homeostatic proliferation in LNs that was similar between sexes **(Extended Data Fig. 11a,b).** However, the majority of naive CD8 T cells during recovery was comprised of CD103^high^ RTEs^75,76^, with a higher prevalence in females **(Extended Data Fig. 11c,d)**. Like in humans, we found significant sex differences in thymus cellularity of mice **(Fig. 5d; Extended Data Fig. 11e,f).** Performing thymectomies (TmX) on young adult mice (**Fig. 5e**) significantly reduced RTEs in LNs (detected by signal joint T cell receptor excision circles, sjTREC), eliminated the sex differences found in middle-aged sham surgery treated animals, and abolished the female advantage in LN cellularity and naive CD8 T cell numbers (**Fig. 5f,g; Extended Data Fig. 12a**). These data mirror the pattern observed in humans, where sex differences are evident when thymic tissue is preserved (score 0–2) but lost upon complete thymic involution (score 3) (**Fig. 5b *right***). Interestingly, disrupting the thymus-derived supply of T cells by TmX in 6 week old mice after the initial establishment of the T cell pool had no impact on the total Tvm cell counts (**Fig. 5h**). These findings are in line with a model in which Tvm cells, largely established from early-life thymic emigrants^77^, persist independently of thymic output. Their stable numbers after TmX in young adults reflect a tightly regulated balance of naive-to-Tvm conversion, shaped by homeostatic constraints such as IL-15 availability and the transient nature of Tvm residency in LNs, as these cells either die or exit the LN, ultimately contributing to LN contraction during aging.

As a result, the loss of naive CD8 T cell replenishment following TmX neutralized sex differences in tumor-specific recognition and CD8 T cell–mediated tumor suppression in middle-aged mice (**Fig. 5i-k; Extended Data Fig. 12b,c**), essentially degrading female anti-cancer immune function in middle-age. Collectively, these results demonstrate that the supply of LNs with de novo generated naive CD8 T cells from the thymus remains relevant even later in life in mice and suggest that thymic output continues to contribute to immune maintenance in middle-aged humans, albeit to a progressively lesser extent in males. Taken together, our data support a two-layer model of naive CD8 T cell loss, in which thymic involution defines the ceiling of T cell supply, while antigen-agnostic Tvm conversion determines how rapidly that supply is eroded.

### Therapeutic thymus regeneration replenishes naive T cells and invigorates tumor-recognition

Thymic atrophy with age results not only from diminished bone marrow–derived lymphoid progenitors and stromal cell senescence, but also from the suppressive effects of sex hormones, particularly androgens^25,78^. While direct evidence is lacking for the extent thymopoiesis in humans can be restored through androgen blockade, clinical data suggest that androgens negatively influences naive CD8 T cell levels^79^. Sex steroid ablation (SSA), which is clinically used to treat hormone sensitive prostate cancer, is currently being evaluated in clinical trials to improve immune regeneration after radiation-induced immunodeficiency^23,80,81^. SSA has shown potential to increase thymopoiesis and TIL presence in melanoma and advanced prostate cancer^5,82,83^. In mice, SSA partially reverses thymic involution and regenerates the T cell compartment in younger animals, but has minimal effect at advanced age^80,84,85^.

To test whether therapeutic SSA could restore naive CD8 T cell availability, improve LN function and drive better tumor recognition, we applied both long-term and short-term strategies in mice: i) surgical gonadectomy (GonX) in early adulthood or middle-age, and ii) pharmacologic SSA using the FDA approved gonadotropin-releasing hormone antagonist Degarelix (DeX) in middle-aged and aged males (**Fig. 6a**). Of note, long-term gonadectomy in females (estrogen depletion via ovariectomy) did not significantly alter thymic cellularity, RTE frequency in LNs (sjTREC content), or naive CD8 T cell numbers (**Fig. 6b,e,f**). In contrast, both surgical and chemical inhibition approaches for androgens led to increased thymus cellularity and normalized thymocyte development, as reflected by restored double-positive (DP) to double-negative (DN) ratios in males **(Fig. 6b,c; Extended Data Fig. 13c,g)**. Importantly, SSA resulted in robust repopulation of LNs, with elevated numbers of RTEs (**Fig. 6d,e**). This translated into a significant restoration of naive CD8 T cell numbers in middle-aged males, recovering from approximately 3x10^5^ cells/ingLN to 8x10^5^cells/ingLN, approaching levels seen in young adults (∼10^6^ cells/ingLN) and untreated middle-aged females. In aged males, DeX treatment also increased naive CD8 T cells, but to a lesser extent, and absolute numbers (2x10^5^ cells/ingLN) remained reduced relative to middle-aged mice (**Fig. 6f; Extended Data Fig. 13a-l**).

**Fig. 6.**
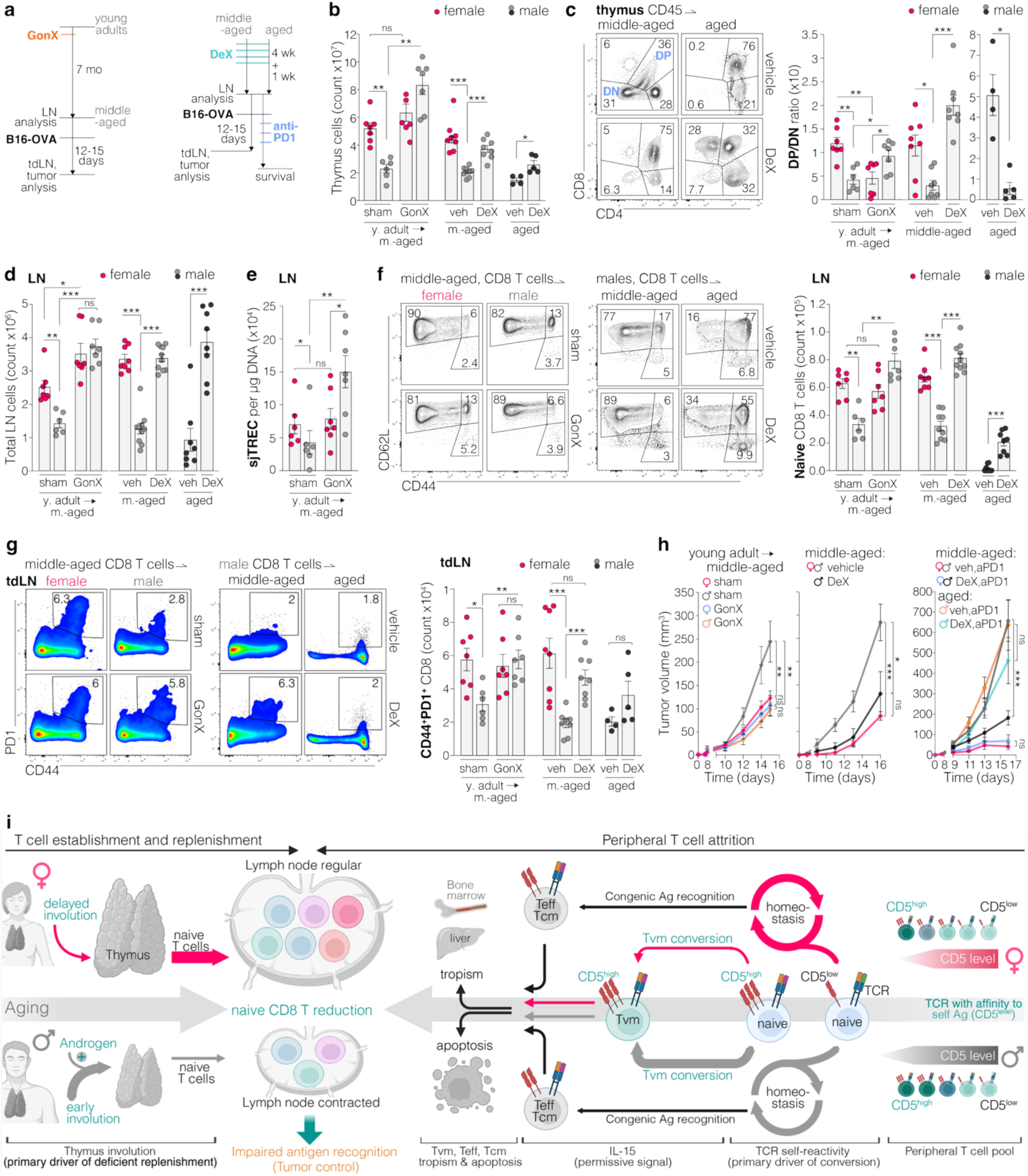
Therapeutic thymus regeneration replenishes naive T cells and reinvigorates tumor-recognition. **a,** Experimental setup, ***left***, young adult and middle-aged mice were gonadectomized (GonX, ovariectomy for females, orchiectomy in males) or, ***right***, middle-aged and aged mice were treated with vehicle (veh) or Degarelix (DeX) before analysis of tumor-naive LNs, or implantation of B16-OVA tumors treated with or without anti-PD1 antibody. **b,** Total count of thymocytes in mice after sham surgery or gonadectomy (GonX) and vehicle (veh) or Degarelix (DeX) treatment in middle-aged and aged mice. **c,** Flow cytometry analysis of thymocytes and double positive (DP) to double negative (DN) ratio**. d,** Inguinal LN cellularity after treatment as described in **a**. **e,** SjTREC quantification in lymphocytes of LNs after sham surgery or GonX. **f,** Flow cytometry analysis of naive CD8 T cells from ingLNs of mice. Gating strategy in **Extended Data Fig. 6f**. **g,** flow cytometry analysis of CD8 T cells from tdLNs. Gating strategy in **Extended Data Fig. 3a**. **h,** B16-OVA tumor growth in, ***left***, sham and GonX (n=6-7) middle-aged mice, ***middle***, vehicle or DeX treated middle-aged mice (n=8/group), and ***right***, vehicle or DeX and anti-PD1 treated middle-aged and aged mice (n=7-13/group). i, Conceptual model linking sex-biased thymic involution and TCR self-reactivity–driven Tvm conversion to naive CD8⁺ T cell loss, LN contraction, and impaired antigen recognition. Data shown as mean ± SEM, each data point representing a single LN or tumor from an individual animal. Statistics calculated with MWU test (b,c,d,e,f,g), two-way repeated measures ANOVA (h) to assess overall effects across time and group; ns=nonsignificant, *p<0.05, **p<0.01, ***p<0.001.

Functionally, these changes eliminated sex-based differences in tumor recognition and response. Following SSA, middle-aged males exhibited increased frequencies of antigen-experienced CD8 T cells in tumor-draining LNs (tdLNs), including CD44⁺PD1⁺ and CD39⁺ subsets, which mark repeatedly stimulated tumor-reactive T cells. Further, tumor-antigen-specific (tetramer⁺) CD8 tumor-infiltrating lymphocytes (TILs) were also increased, mirroring responses seen in females. Correspondingly, tumor growth dynamics equalized between sexes (**Fig. 6g,h; Extended Data Fig. 14a-i**), with SSA treated males controlling tumor growth better than control treated male mice. Preconditioning of middle-aged mice with DeX restored numbers of naive CD8 T cells in LNs and increased the therapeutic response to subsequent anti-PD1 immunotherapy (**Fig. 6h *right*; Extended Data Fig. 14j**).

Together, these findings demonstrate that SSA can reinvigorate thymic output, replenish the naive CD8 T cell pool, and enhance antitumor immunity in middle-aged males, thereby eliminating functional sex differences. These data also show that improving naive CD8 T cell pools in middle-aged males can increase responses to anti-PD1 immunotherapy. However, the diminished efficacy of this intervention in aged males highlights the narrowing therapeutic window for immune regeneration later in life.

## Discussion

Our study highlights substantial sex- and age-related disparities in naive CD8 T cell homeostasis and its connection to functional antigen recognition in LNs, illuminating a previously underappreciated link between naive T cell dynamics, thymic involution and impaired cancer immunity during aging. Age-related thymic involution and its consequences for peripheral T cell homeostasis have been studied for decades, with foundational work establishing that progressive loss of thymic output leads to numerical and functional decline of naive T cells and contraction of TCR diversity with age^19–21^. More recent clinical evidence has linked thymic atrophy directly to increased cancer risk and mortality in humans^20,21,86^, elevating this process—once-considered passive—to an active determinant of immune competence. We demonstrate that males experience an earlier and more profound decline in naive CD8 T cell abundance associated with a contraction of LNs that critically limits the number of diverse clones necessary for effective antigen recognition. This deficiency in males results in a skewed intratumoral T cell composition characterized by an increased frequency of antigen-naive bystander T cells and fewer activated tumor-reactive effectors, which compromises anti-cancer immunity in middle age. Although T cell infiltration in tumors generally correlates positively with prognosis, the functional status and antigen specificity of TILs are paramount determinants of clinical outcomes. Our findings align with reports demonstrating that increased frequencies of naive-like TILs, often associated with age^7^, correlate with ineffective anti-cancer responses and poor therapeutic outcomes, highlighting the need for functional rather than numerical assessments of TIL populations in clinical contexts ^29,31–34^. While aged mice exhibit a significant reduction in TCR diversity of up to 80%, which leads to measurable deficits in immune functionality, it remains less clear whether a reduction in T cell clones has equivalent consequences in humans^87–90^. Notably, LNs in aged humans maintain relative clonal diversity^91^. However, our data suggest that a functional T cell pool depends not only on the relative diversity but also on the absolute number of unique T cell clones available in draining LNs. Recent studies^15,17^ have shown that tdLNs continuously generate stem-like CD8 T cell populations that serve as a renewable source of effector cells and are enriched for high-affinity clones. In this context, the size and diversity of the naive T cell pool represent a key upstream determinant of the landscape from which these stem-like CD8 T cell populations are derived. Thus, reduced naive T cell availability can constrain anti-tumor immunity by limiting the generation of high-affinity, stem-like T cell clones capable of sustaining effective responses. Importantly, this mechanism reflects a fundamental relationship between naive CD8 T cell availability in LNs and the stochastic probability of antigen recognition. In this framework, the tumor serves primarily as a source of locoregional antigen, while the LN naive T cell pool determines the likelihood that a cognate clone will be recruited^17^. Tumor immunogenicity acts as a modulating variable in this relationship, as a higher density of neo-antigens and antigens with specific recognition motifs^92^ can partially compensate for a reduced naive precursor pool, demonstrated in the context of poorly versus highly immunogenic tumors^93^. In support of the broader applicability of this framework, age-related impairment of tumor control has been documented across a range of syngeneic tumor models with and without immunogenic modifications and consistently implicate loss of tumor-specific CD8 T cell responses as a shared underlying mechanism^7,8^.

A central insight of our study is the dual mechanisms driving naive CD8 T cell loss in males: an accelerated, antigen-agnostic conversion into Tvm cells combined with a faster decline in thymic output **(Fig. 6i)**. Prior work has shown that total peripheral T cell numbers, largely driven by the CD4 compartment, remain relatively stable with age through homeostatic proliferation and increased cell lifespan^50^, whereas CD8 T cells progressively decline^41–44^. In line with a recent publication^20^, our data refine this paradigm by demonstrating a selective depletion of naive CD8 T cells, driven in part by premature differentiation into Tvm cells. This loss is insufficiently compensated in males due to earlier thymic involution. Peripheral Tvm conversion is what makes declining thymic output consequential. Without ongoing attrition, the long lifespan of naive T cells would buffer against even substantially reduced thymic replenishment, and the sex-biased divergence we observe would not emerge on the timescale of middle age. The sex-biased reduction in naive CD8 T cells profoundly reduces the numerical abundance of unique clones, critically impairing antigen recognition capacity in tumor-draining LNs. Human data **(Fig. 2g,h**; **Fig. 5a-c; Extended Data Fig. 5d; Extended Data Fig. 10a-e)** further support this model, showing that while RTEs decline in both CD4 and CD8 compartments with age, only naive CD8 T cells undergo a marked numerical reduction. This likely reflects a vulnerability of CD8 T cells to antigen-agnostic differentiation into Tvm cells, a process that is not a characteristic feature of naive CD4 T cells. CD8 Tvm formation depends on two features: clone-specific self-reactivity and responsiveness to homeostatic cytokines such as IL-15. Because MHC class I-restricted CD8 T cells sample a broader and more continuous range of endogenous self-peptides than CD4 T cells, CD8 T cells may display greater heterogeneity in their baseline self-reactivity^18^. CD4 T cells exhibit a lower propensity to lose their naivety through antigen-agnostic memory differentiation in mice^51,94,95^ and are less susceptible to phenomena like excessive memory formation, which occurs with CD8 T cells during chronic infections with cytomegalovirus or Epstein-Barr virus in humans^96^. Consequently, while the systemic population of CD4 T cells remains relatively stable over time^50^, independent of age and sex, naive CD8 T cells are more prone to Tvm formation, senescence and apoptosis, suggesting that maintaining an abundant and clonally diverse pool of naive CD8 T cells becomes increasingly dependent on de novo thymic synthesis with age. An additional layer of complexity arises from recent work showing that the Tvm compartment is heterogeneous and includes a Ly49⁺CD122⁺ CD8 regulatory-like subset^97,98^. Although we did not delineate this population in our study, they might contribute to impaired tumor control. However, the long-term impairment we observe is consistent with persistent contraction of the naive CD8 T cell pool with age. Future studies will be required to determine whether regulatory Tvm subsets are preferentially expanded under these conditions and how they influence tumor immunity.

Our results support the critical role of thymic output in maintaining naive CD8 T cell populations in adulthood, echoing recent clinical findings linking thymic atrophy with increased cancer risk and mortality in humans^18^. The observed sex disparity in thymic involution during the transition from young adulthood to middle age suggests a hormonal influence, with androgens exerting potent inhibitory effects on thymopoiesis^25^. Augmenting lymphoid progenitor cells in the bone marrow by depleting competing myeloid-biased hematopoietic stem cells has shown promise in improving adaptive immune responses but results in only modest increases in naive T cells^99^, presumably due to the lack of thymus-based T cell maturation in advanced age. In our study, therapeutic thymus regeneration via androgen blockade restored naive CD8 T cell numbers, enhanced cancer antigen recognition, and reversed sex-related differences in cancer immunity in middle-aged male mice **(Fig. 6)**. These data underscore the potential of therapeutic interventions targeting thymic regeneration, especially within a potential therapeutic window before advanced age-related changes such as thymus epithelial cell senescence, LN stromal fibrosis and structural degeneration irreversibly hinder regeneration efforts^85,100,101^.

Beyond strategies aimed at enhancing thymic output, limiting premature Tvm differentiation may represent a complementary way to preserve naive CD8 T cell numbers. Our finding that transient artificial induction of Tvm cells in females recapitulates male-like aging phenotypes suggests that early-life skewing toward Tvm can have durable effects on immune homeostasis. Mechanistically, Tvm formation depends on IL-15 availability and clone-specific self-reactivity^51,52^, but how these factors are regulated in physiological settings remains incompletely understood. Recent work has also identified thymic precursors with virtual-memory–like features^54^, indicating that developmental biases may contribute to baseline Tvm abundance. Rather than implying direct therapeutic tractability, our data highlight the need to define the upstream checkpoints that govern Tvm differentiation across both thymic and peripheral compartments, and to determine whether modulating these pathways could help delay naive CD8 T cell depletion during aging.

Clinically, our findings highlight a potential but context-dependent role for naive CD8 T cell availability in shaping anti-tumor immunity. Immune checkpoint blockade (ICB) does not universally depend on de novo T cell priming, and age alone is not a reliable predictor of therapeutic outcome in patients^102–104^. Nevertheless, in settings where antitumor responses require recruitment of new T cell clones, such as in early-stage disease, neoadjuvant treatment or tumors with limited pre-existing immunity, the numerical availability of naive CD8 T cells in draining LNs may influence treatment efficacy. Our mouse studies show that transient restoration of thymic output expanded the naive CD8 T cell pool in middle-aged male mice and improved responses to anti-PD-1 therapy in a model system where priming is rate-limiting **(Fig. 6h)**. These data do not imply that naive T cell decline is a dominant determinant of ICB efficacy in all patients but rather demonstrate that enhancing thymic function can augment anti-tumor immunity when naive clone recruitment is limiting.

The human T cell and thymus data in our study are derived from at least two independent cohorts, strengthening the translational relevance of our findings. The CT-based LN volume data are inherently more exploratory. The scans were obtained from individuals with unknown health indications, volumes are unadjusted for body composition or underlying conditions, and LN size and number can vary substantially between individual humans compared to mice. Nevertheless, averaging volumetric measurements across anatomically defined LN regions and using three-dimensional volume rendering show the sex- and age-related trends we observe are consistent with our murine and human immune cell data. These convergent lines of evidence raise the possibility that thymus-targeted interventions, such as short-term SSA treatment, could be explored in selected clinical contexts where insufficient priming constrains immunotherapy responses.

Collectively, our study provides compelling evidence for the intertwined roles of naive T cell dynamics and thymic functionality in shaping sex- and age-specific LN function and immune competence. Future research should focus on dissecting molecular pathways governing thymic involution and regeneration—identifying targets for intervention that do not rely solely on hormone ablation—and on developing personalized strategies accounting for individual hormonal profiles and age-related immune trajectories. Ultimately, enhancing thymic function and naive T cell resilience holds significant promise for improving immune responsiveness against cancer and infections, thus promoting healthier aging.

## Online methods

### Mice

Wild type C57BL/6 mice, *Lta*^-/-^ (lymphotoxin alpha knockout mice B6.129s2-Ltatm1Dch/J from the Jackson Laboratory, strain #002258), CD45.1 congenic mice (B6. SJL-PtprcaPepcb/BoyJ, from Jackson Laboratory, strain #002014) and OT-1 (C57BL/6-Tg(TcraTcrb)1100Mjb/J) were used for animal models. All mice were bred and maintained in animal facilities of the Massachusetts General Hospital with housing in accordance with procedures outlined in the National Institutes of Health’s Guide for the Care and Use of Laboratory Animals. Mice received a standard pelleted rodent diet and tap water ad libitum and were maintained at 20-24°C, 70 % humidity and 12/12-hour light/dark cycle. All experiments were performed at the Massachusetts General Hospital (MGH) and approved by the Institutional Animal Care and Use Committee (IACUC) of the MGH under protocols 2023N000101, 2011N000085 and 2004N000123. Age- and sex-matched groups of adolescents (4-5 weeks), young adults (7-10 weeks), middle-aged (9-12 month) and aged (>18 month-old) female and male mice were used in each experiment unless specified otherwise.

### Animal models and tumor cell lines

The B16F10 melanoma cell line (ATCC) was cultured in DMEM supplemented with 10% heat-inactivated fetal bovine serum (FBS, Gibco), 1% penicillin–streptomycin (Gibco), and 1% L-glutamine (Gibco) at 37°C and 5% CO₂. B16-OVA melanoma cells (ovalbumin-expressing B16F10) were kindly provided by Dr. Susan Thomas (Georgia Institute of Technology). B16-SIINFEKL-zsGreen cells were generated in-house as previously described^105^. To exclude sex-mismatched antigen effects, B16F10 cells were verified to lack Y-chromosome–encoded sequences, ensuring that tumor recognition was not influenced by sex-specific histocompatibility antigens (see “PCR to determine sex from genomic DNA” in methods) **(Extended Data Fig. 2a)**. All cell lines were used at low passage number (below passage 15) and regularly tested for mycoplasma contamination using the MycoAlert Mycoplasma Detection Kit (Lonza). Only mycoplasma-negative cultures were used for in vivo experiments. For tumor engraftment, B16F10 or B16-OVA cells (5×10^5^) were injected intradermally into the lower flank of mice. Animals were monitored daily, and tumor volumes were measured every 2–3 days. Tumor volume was estimated as: tumor volume = width × length × (width/2). For survival analyses, animals were euthanized based on approved humane endpoints once the tumor exceeded 1,000 mm³ or reached 2 cm in any diameter.

### Study population

The Swedish CArdioPulmonary bioImage Study (SCAPIS) is a general population-based prospective study^71^. Middle-aged participants (50–64 years old females and males) underwent comprehensive clinical examination including imaging and functional studies of the heart and lungs—that included the thymus—and completed extensive questionnaires to assess lifestyle as well as prior and current medication regiments. The SCAPIS Leukocyte subcohort of 1078 participants was consecutively recruited from the SCAPIS center at Linköping University Hospital to establish an immune cell profile in whole fresh blood using flow cytometry^73^. Twenty-nine participants (2.7%) were excluded from analysis: 9 (0.8%) due to missing images, 6 (0.6%) with anterior mediastinal lymphadenopathy or tumor, 5 (0.5%) with prior surgery preventing evaluation, 2 (0.2%) with imaging artefacts, and 7 (0.6%) for miscellaneous anatomical or technical reasons.

### Volume segmentation in CT scans

Computed tomography (CT) scans of human individuals are from the Cancer Imaging Archive of the National Institutes of Health (NIH)^38,39^ and were analyzed using 3D Slicer (version 5.6.2). These datasets do not include clinical data, and thus the health status of the individuals is unknown. To ensure that LN measurements reflected non-pathologic anatomical variation, we excluded all scans showing radiologic evidence of lymphadenopathy or other abnormalities that could influence LN size.

#### Lymph node

(LN) analysis: Body scans were cropped to the axillary or inguinal regions. Superficial skin areas were excluded. The segmentation threshold was adjusted to include LNs while excluding surrounding fatty tissue and skin. Using 3D volume rendering, LNs were identified and surrounding tissues removed manually until only individual LNs remained. The average LN volume per individual was used for statistical analysis. Datasets with one or more pathologically enlarged LNs (>700 mm³) were excluded.

#### Spleen

analysis: Scans were cropped to the region beneath the lungs to isolate the spleen. The segmentation threshold was adjusted to include the spleen while excluding surrounding fatty tissue. 3D volume rendering was used to identify the spleen and manually remove extraneous tissue.

#### Thorax

analysis: Scans were cropped between the clavicles and the upper pelvis. The segmentation threshold was set to include all tissue within the thorax. Using 3D volume rendering, objects not part of the human thorax (e.g., scanning table) were removed.

#### Thymus

analysis: scans were assessed as described^73^. In brief, thymus fat and soft tissue content was visually analyzed and graded on a scale 0–3, grade score 0: predominantly soft tissue, score 1: parity of fatty and soft tissue; score 2: predominantly fat; score 3: complete fatty replacement **(Extended Data Fig. 9a)**.

#### Thymus analysis of the SCAPIS cohort

CT scans of the thorax was performed along the guidelines of the SCAPIS protocols ^71^. All scans were reviewed using a fixed window setting and evaluated by an experienced thoracic radiologist ^73^.

### Organ resection procedures

All surgical procedures were performed under Ketamine/Xylazine (ip) anesthesia. Procedures were performed under a dissecting microscope where applicable. Animals were monitored daily post-surgery for signs of pain, infection, abnormal behavior, and body weight.

#### LN dissection

A 0.5 cm skin incision exposed the inguinal LN. Vessels were compressed with fine curved forceps, and the LN was excised. The skin was closed with wound clips, and mice recovered for ≥14 days.

#### Splenectomy

The spleen was accessed via an incision through the skin and peritoneum at the right flank. Vascular connections at the splenic hilum were ligated, and the spleen was removed. The peritoneum and skin were closed, and mice recovered for ≥14 days.

#### Thymectomy

A semicircular skin incision along the upper thorax was followed by a partial sternal incision to expose the anterior mediastinum. Both thymic lobes were identified and removed by blunt dissection, taking care to avoid injury to blood vessels and pericardium. Any remaining thymic tissue was removed by gentle aspiration. The chest wall was closed with absorbable suture and the skin with wound clips. Mice were allowed to recover and mature to middle age before experiments. Complete thymic removal and absence of thymic regeneration were verified visually during surgery and confirmed at experimental endpoint.

#### Orchiectomy

Mice were placed in dorsal recumbency and the scrotal region was shaved and disinfected. The testis, epididymis, and associated fat pad were gently exteriorized through a small scrotal incision. A ligature of absorbable suture was placed below the testis and epididymis, which were then excised. The scrotal incision was closed with absorbable suture. Successful orchiectomy was confirmed by measurement of serum testosterone levels by ELISA at experimental endpoint **(Extended Data Fig. 16b,f)**. Mice recovered for ≥14 days before downstream analysis.

#### Ovariectomy

Mice were placed in ventral recumbency and the dorsal skin overlying the ovaries was shaved and disinfected. Ovaries were accessed through bilateral midline incisions along the back. A ligature of absorbable suture was placed at the oviduct-uterus boundary, and the ovaries were excised. The peritoneum was closed with absorbable suture and the skin with wound clips. Successful ovariectomy was confirmed by atrophy of the uterine horns at experimental endpoint. Mice recovered for ≥14 days before downstream analysis.

### Chemical sex steroid ablation

Degarelix (MedChemExpress, Cat. #214766-78-6) was dissolved in a solution of 10% DMSO, 40% PEG3000, 5% Tween-80, and 45% saline. It was administered subcutaneously at 0.25 mg once weekly for four consecutive weeks. Control mice received vehicle only.

### In vivo treatments

Mice were injected intraperitoneally with immune-modulating antibodies or cytokine complexes. For PD-1 blockade, anti-PD-1 mAb (clone RMP1-14, Bio X Cell) was administered at 10 mg/kg three times every 3 days. For T cell depletion, anti-TCRβ mAb (clone H57-597, Bio X Cell) was given at 10 mg/kg once. For CD8 T cell depletion, anti-CD8α mAb (clone 53-6.7, Bio X Cell) was administered at 10 mg/kg, four times every 5 days. Efficient CD8 T cell depletion was confirmed by flow cytometric analysis of peripheral blood before continuation of downstream experiments **(Extended Data Fig. 2c)**. For IL-15/IL-15Rα complex (IL-15 Cx) treatment, IL-15 Cx (6 µg, Sino Biological, cat. #CT113-M08H) or vehicle (0.9% NaCl solution) were administered once per week for 4 times.

### Immunization of mice

Tumor cell lysates were prepared by subjecting 1×10^8^ cells/mL in sterile water to five freeze-thaw cycles (liquid nitrogen and 37°C water). Fifty microliters of tumor cell lysate or DQ-OVA solution (Invitrogen, cat. #D12053) containing 10 µg of LPS (Sigma-Aldrich, cat. #L4774) as adjuvant, or PR8 virus particles (1×10^5^ inactivated particles with 2 µg poly(I:C) and 2 µg anti-CD40 mAb [clone MR-1, Bio X Cell]) or respective controls were injected intradermally into the lower flank. Anti-CD40 antibody was included to mimic CD4 T cell help and promote efficient DC activation and cross-priming. Draining inguinal LNs and the contralateral inguinal (non-draining) LNs were analyzed 24- or 72-hours post-immunization.

### Tissue Dissection and single-cell preparation

Mice were euthanized, and tissues were resected and mechanically dissociated. Individual LNs were mechanically dissociated by gently pressing them with a syringe plunger against the surface of a sterile petri dish. Single-cell suspensions from the thymus were filtered through a 40 µm strainer. Single-cell suspensions from tumor and spleen were filtered through a 70 µm strainer. Cells in the resulting suspension were counted before any filtering or centrifugation to ensure reliable cell counts without technical artifacts. For flow cytometry analysis of antigen presenting cells, LNs were treated with 1 mg/ml Collagenase-D (Sigma) and 0.1 mg/ml DNAse-I (Roche, in HBSS) for 30 min at 37°C before mechanical dissociation. Blood and spleen samples underwent hemolysis with Ammonium-Chloride-Potassium (ACK) buffer (Fisher Scientific, cat. #50-983-219).

### Flow cytometry

Single-cell suspensions were incubated with LIVE/DEAD Zombie dye (BioLegend, cat. #423108) and anti-mouse CD16/32 in PBS for 15 min at 4°C to block Fc receptors and stain for viability. When applicable, tetramers were added for 1 hour at room temperature. Surface staining was then performed with fluorochrome-conjugated antibodies in flow buffer (2% FCS, 0.5 mM EDTA in PBS) for 20 min at room temperature (antibody list in **Extended Data table 1**). After washing, cells were fixed and permeabilized using the Foxp3/Transcription Factor Staining Set (eBioscience, cat. #00-5523-00) for 30–60 min at 4°C, followed by intracellular staining for ≥1 hour at 4°C. Data were acquired using an Aurora spectral flow cytometer (Cytek, 5 laser configuration) and analyzed with FlowJo (v.10.1, TreeStar). Gating strategies are shown for murine lymphocytes (**Extended Data Fig. 6f**), murine tumor samples (**Extended Data Fig. 3a**), murine thymocytes (**Extended Data Fig. 11e**), and human PBMCs (**Extended Data Fig. 5g**).

### In Vitro Stimulation and Proliferation Assays

Sorted CD8 naive T cells were stained with carboxyfluorescein succinimidyl ester (CFSE) per manufacturers’ instructions (Thermo Fisher, cat. #C34554) or left unstained, then stimulated on plates coated with anti-CD3 (5 µg/ml, Biolegend, cat. #100340) and anti-CD28 (4 µg/ml, Biolegend, cat. #102116) antibodies. The cells were incubated for 48 hours in complete RPMI1640 medium supplemented with 10% FBS (Gibco), 1% penicillin/ streptomycin (Gibco), 1% l-glutamine (Gibco), 1% HEPES (Gibco) and 0.1% β-mercaptoethanol (Sigma) at a concentration of 5x10^5^ cells per well in a 96-well flat-bottom anti-CD3/anti-CD28 coated plate. For the last 4 hours of incubation Brefeldin-A (Biolegend, cat. #420601) was added. Intracellular staining for IFNg, IL-2, and TNF was performed as described above.

### mRNA Sequencing and Analysis

Naive CD8 T cells were defined as CD44⁻CD49d⁻CD122^-^ and virtual memory (Tvm) cells as CD44⁺CD49d⁻CD122⁺. Cell sorting was performed on a Sony MA900 cell sorter, and a minimum of 200,000 cells per population were collected. 100,000 sorted cells were immediately resuspended in 50 µL Zymo DNA/RNA Shield and stored at RT until further processing. The remaining 100,000 cells per sample were activated on anti-CD3/CD28 coated 96-well plates (Ultra LEAF™ Purified anti mouse CD3ε, 5ug/mL, Ultra LEAF™ Purified anti mouse CD28 4ug/mL) and incubated for 5 hours before resuspending in 50 µL Zymo DNA/RNA Shield. Three biological replicates were collected per group. Total RNA extraction, library preparation, sequencing and data processing were performed by Plasmidsaurus (Eugene, OR). Briefly, total RNA was extracted from sorted cells preserved in Zymo DNA/RNA Shield using a bead-based extraction approach. RNA concentration was assessed by fluorescence-based quantification prior to library preparation. Sequencing libraries were generated using a 3′-end counting approach: mRNA was converted to cDNA via reverse transcription using a poly(dT)VN primer, followed by second-strand synthesis, tagmentation, indexing, and amplification. Unique molecular identifiers (UMIs) and unique dual indices (UDIs) were incorporated to enable PCR deduplication and prevent index hopping. Single-end sequencing (∼90 bp read length) was performed on an Illumina platform. Raw FASTQ files were generated and demultiplexed using BCL Convert (v4.3.6) and fqtk (v0.3.1). Read quality filtering was performed with FastP (v0.24.0), including poly-X tail trimming, 3′ quality-based trimming, a minimum Phred score of 15, and a minimum read length of 50 bp. Filtered reads were aligned to the mouse reference genome (GRCm38/mm10) using STAR (v2.7.*) with non-canonical splice junction removal. BAM files were coordinate-sorted with samtools (v1.21), and PCR/optical duplicates were removed by UMI-based deduplication using UMICollapse (v1.1.0). Alignment quality metrics were assessed with RustQC (v0.2.1) and summarized using MultiQC (v1.33). Gene-level counts were quantified with featureCounts (Subread v2.1.1) using strand-specific counting, with exons and 3′ UTRs as feature identifiers grouped by gene_id. Sample-level quality and reproducibility were assessed by principal component analysis (PCA) and pairwise Pearson correlation of TMM-normalized counts. Differential gene expression analysis was performed using edgePython (v0.2.5). Genes with a false discovery rate (FDR) < 0.05 and an absolute log₂ fold change > 1 were considered differentially expressed. Genes used for senescence genes sets, cell cycle arrest: *Cdkn1a, Cdkn2a, Cdkn1b, Cdkn2b, Rb1, Trp53*; inflammatory secretion/SASP: *Il6, Il1a, Il1b, Cxcl1, Ccl2, Ccl5, Tnf, Igfbp3, Igfbp7, Mmp3, Mmp9*; DNA damage response: *H2ax, Atr, Atm, Chek1, Chek2, Brca1, Trp53bp1*; Resistance to apoptosis: *Bcl2, Bcl2l1, Mcl1, Birc5, Cflar*; Oxidative stress: *Nox4, Hmox1, Sod1, Sod2, Cat, Gpx1*; Terminal differentiation: *Klrc1, Klrc2, Klrk1, Klrb1c, Klrd1*.

### Immunohistochemistry

Tissues were fixed in 4% paraformaldehyde (PFA) at 4°C overnight (ON) and incubated in 30% sucrose (Sigma-Aldrich) ON at 4°C. Samples were embedded in OCT compound (Tissue-tek), frozen, and cryosectioned (10 µm). Sections were washed in PBS, blocked (5% goat serum, 0.25% Triton X-100 in PBS) for 1 h at room temperature, and incubated with primary antibodies overnight at 4°C. After washing, sections were incubated with secondary antibodies (Jackson ImmunoResearch), counterstained with 4′,6-diamidino-2-phenylindole (DAPI), and mounted in VECTASHIELD PLUS Antifade Mounting Medium (Vector Laboratories, cat. #H-1900-2). Images were acquired using an AxioScan Z.1 microscope (Zeiss) and analyzed with ImageJ.

### Enzyme-linked immunoassay

Serum ***testosterone*** levels were measured using a testosterone ELISA kit (Crystal Chem, cat. #80552) following the manufacturer’s instructions.

For ***IL-15/IL-15Rα complex*** (IL-15 Cx) measurements, individual inguinal LNs were weighed and homogenized in 100 µL radioimmunoprecipitation assay (RIPA) buffer. Fifty microliters of lysate per sample were used for ELISA (Thermo Fisher Scientific, cat. #BMS6023) according to the manufacturer’s protocol.

### PCR to determine sex from genomic DNA

Genetic sex of tumor cell lines was determined as described^106^. DNA was extracted (DNA extraction columns from Zymo Research) from cultured tumor cells (B16F10, female controls: 4T1, Yumm3.3) or male splenocytes. PCR was performed using primer pairs listed in **Extended Data table 2** with Hot Start Taq Plus polymerase (Qiagen, cat. #203605). Parameters: 95°C for 2 min, 35 cycles with 94°C for 30 s, 56°C (*SX* and *Ube* primers) and 55°C (*Zfy* primers) for 30 s, and 72°C for 30 s, followed by final elongation at 72°C for 5 min. PCR products were resolved on 2% agarose gels and visualized with GelRed (Biotium).

### sjTREC PCR detection in mice

Genomic DNA from LN cells was extracted using the DNeasy Blood & Tissue Kit (Zymo Research) per the manufacturer’s instructions. qPCR with primers and probe specific for single joint T cell receptor excision circles (sjTREC, **Extended Data table 2**) was performed using PrimeTime™ Gene Expression Master Mix (IDT, cat. #1055770). Parameters: 95°C for 3 min, 45 cycles of 95°C for 15 s, 60°C for 60 s, followed by final elongation at 72°C for 5 min. Standard curves were generated using a plasmid containing the sjTREC sequence (gift from Karin Gustafsson, MGH). sjTREC copy number per µg DNA was calculated from the standard curve.

### TCR sequencing

CD8 T cells were isolated from individual inguinal LNs using the EasySep™ Mouse CD8 T Cell Isolation Kit (StemCell, cat. #19753). Sorted cells were submitted to iRepertoire for TCR sequencing and bioinformatic analysis. This included RNA isolation, reverse transcription, amplification of *TRA* and *TRB* CDR3 regions, and sequencing to assess T cell clonality and diversity.

#### Shannon entropy

Shannon entropy was calculated to determine TCR diversity. The frequency of each clonotype was used to compute probabilities (*p*_i_ = frequency_i/*N*_total), and entropy (*H*) was calculated as:

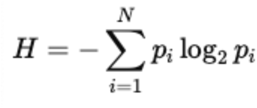

where *N* is the total number of unique clonotypes.

#### Shannon entropy approximations

The average values of total TRB and unique TRB of all 6 samples (3 female, 3 male) were used to calculate theoretical values of samples with similar characteristics and a certain diversity (100% and 50% unique TCRs). Entropy was computed using the formula above). Probabilities for unique clonotypes were calculated as *p*_unique_=1/*N*_total_, while probabilities for expanded clonotypes were based on their observed frequencies. The total entropy was obtained by summing the contributions from unique and expanded clonotypes.

(Case **A**, 100 %) maximum diversity means each sequence is unique and appears exactly once, calculation with *H*=log_2_*N*;

(Case **B**, 50 % with a single highly expanded clonotype):

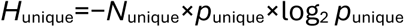

equation for expanded clonotypes: *H*_expanded_=−*p*_expanded_×log_2_ *p*_expanded_

equation for combined entropy: H_total_=H_unique_+ H_expanded_

(Case **C**, 50 % duplicates evenly distribution among 10 expanded clonotypes):

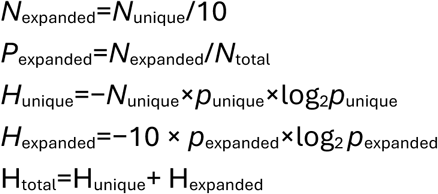

### Re-analysis of spectral flow data

Spectral flow data from Bohacova *et al.*^47^ (Synapse: syn53238646) were analyzed using the gating strategy as described in **Extended Data Fig. 5f** with the aim to quantify true naive CD8 and CD4 cells and with an emphasis to exclude all non-true naive T cells that might interfere with the subsequent quantification of recent thymic emigrants (RTEs)^47^. The following markers were used in **live cells**: CXCR5^-^/CD19^-^ (excludes B cells, T follicular helper cells), CD14^-^/CD123^-^ (excludes myeloid cells), NKp80^-^/CD56^-^ (excludes NK cells), TCR γ/ δ ^-^/CD3^+^ (excludes γ/δ T cells, include all other T cells), gating on **CD8 T cells**: CD25^-^/PD1^-^/FAS^-^/GZMB^-^ (excludes cells expressing activation markers), CD27+/CD45RA+ (true naive CD8 T cells), CD38^high^ (RTE). **CD4 T cells**: FAS^-^/CXCR3^-^ (excludes cells expressing activation markers), CD127^+^/CD25^-^ (excludes regulatory T cells), CD27^+^/CD45RA^+^ (true naive CD8 T cells), CD38^high^ (RTE). Data were analyzed with FlowJo (v.10.1, TreeStar).

### Re-analysis of scRNA-seq data

All scRNA-seq data were analyzed with Seurat (v.5).

#### Human PBMC

public scRNA-seq data^46^ from Terekhova et al.^46^ Synapse: syn49637038, were filtered for CD8 T cells and assigned a “*naive score*” using the Ucell package with positive and negative modulator genes (*LEF1^+^, TCF7^+^, KLF2^+^, BACH2^+^, FOXP1^+^, GZMB^-^, PRDM1^-^, TBX21^-^, FAS^-^, EOMES^-^*).

#### Human LN cells

from public scRNA-seq data^48^, were filtered for CD8 T cells and assigned a “*naive score*” as described above. Antigen presenting cells (APCs) from the same data set were analyzed using the cell subset attribution from the original publication. scTCR-seq data of T cells from LNs of the same data set were analyzed using the scRepertoire^107^ (v.2.3.0) package in R.

#### Human melanoma

data sets from Durante *et el.*^27^ GSE139829; Sade-Feldman *et al.*^28^ GSE120575; Li et al.^29^ GSE123139; Zhang et al.^30^ GSE215121). Cells of individual patients were filtered for CD8 T cells and the percentage of naive, cytotoxic and exhausted CD8 T cells were attributed with module scores using the Ucell package (naive: *SELL, CCR7*; cytotoxic: *GZMA, GZMB, PRF1, IFNG, PTPN6, GNLY*; exhausted: *HAVCR2, TOX, TIGIT*).

#### Human thymocyte

public scRNA-seq data from Park et al.^60^ (https://zenodo.org/records/3711134) and Cordes et al.^61^ GSE195812, were assigned for subsets using the Ucell package with positive and negative modulator genes (double negative (DN)1: *CD34^+^, HES1^+^, IL7R^+^, TRAC^-^;* DN2: *HES1^+^, PTCRA^+^, TRAC^-^*; *DN3: NOTC1^+^, PTCRA^+^, CD8A^-^, CD4^-^;* DN4: *RAG1^+^, BCL11B^+^, PTCRA^+^, CD8A^-^, CD4^-^*; double positive (DP): *RAG1^+^, CD8A^+^, CD4^+^*; single positive (SP) CD8: *CD8A^+^, CD8B^+^, TRAC^+^, CD4^-^*; SP CD4: *TRAC^+^, CD4^+^*; Treg: *FOXP3^+^, IL2RA^+^, IKZF2^+^*; mature naive: *CCR7^+^, TRAC^+^, FOXP1^+^, KLF2^+^, CD8A^+^, CD4^+^*; *CD69*^-^, *IL2RA*^-^, γδ T cells: *TRDC^+^, SOX13^+^, TRAC^-^*; *CD8A*^-^, *CD8B*^-^, *CD4*^-^.

### Immune scores

Immune scores were derived from transcriptomic data of melanoma tumors from The Cancer Genome Atlas (TCGA) using the ESTIMATE (Estimation of Stromal and Immune cells in Malignant Tumor tissues using Expression data) platform developed at MD Anderson Cancer Center, University of Texas.

### Statistical analysis

Groups were compared using Prism software (GraphPad, v.10). The statistical methods used are described in the figure legends and include Mann-Whitney U test (MWU), independent-sample t test, log-rank test, 2-way ANOVA, Fisher’s exact test and Pearson correlation. Tumor growth curves were analyzed using two-way repeated measures ANOVA with Geisser-Greenhouse correction, with the interaction between time and group used to assess differences in growth dynamics between experimental groups. Multiple pairwise comparisons were not performed unless explicitly indicated. In all mouse experiments, each data point represents a single LN or tumor from an individual animal. In the SCAPIS data, a sex effect controlling for age was evaluated using General Linear Model (GLM). Statistical significance is indicated in figures as ns=nonsignificant, *p<0.05, **p<0.01, ***p<0.001.

## SUPPLEMENTAL MATERIALS

Extended Data Fig. 1 to 14

Extended Data table 1 and 2

Supplementary Data Fig. 1 to 2

## Supporting information

Supplementary Informations

## Acknowledgments

We thank Dr. Peigen Huang and the Cox-7 animal facility for maintaining and providing mice and technical support. We thank Mark Duquette for providing help and technical assistance. We thank Dr. Marina Terekhova, Dr. Pavla Bohacova and the lab of Dr. Maxim N. Artyomov for granting access to data sets of human PBMCs. We thank Dr. David Scadden and Dr. Karin Gustafsson for critical discussions and experimental support. The results shown in Extended Data Fig. 1c are in whole or part based upon data generated by the TCGA Research Network: https://www.cancer.gov/tcga. We thank the NIH Tetramer Core Facility (contract number 75N93020D00005) for providing tetramers for flow cytometry.

## Funding

German Research Foundation grant ME 5486/1-2 (LM)

National Institutes of Health grant F32CA275298 (MJO)

National Institutes of Health grant K00CA234940 (HZ)

National Institutes of Health grant R21AG072205 (TPP)

National Institutes of Health grant R01CA214913 (TPP)

National Institutes of Health grant R01HL128168 (TPP)

National Institutes of Health grant R01CA284372 (TPP)

National Institutes of Health grant R01CA284603 (TPP, LLM)

The Rullo Family MGH Research Scholar Award (TPP)

Melanoma Research Alliance (DRS)

V Foundation for Cancer Research (DRS)

NIAID DP2 AI176139 (DRS)

NIAID U19 AI082630 (DRS)

Heart–Lung Foundation, Sweden (20180436)

Swedish Research Council (2018–03232)

## Author contributions

Conceptualization: LM, TPP

Methodology: LM, TPP, DRS, MZ, LLM

Investigation: LM, MZ, MJO, HZ, LL, PL, MS, LJ

Visualization: LM, LLM

Supervision: TPP

Writing the original draft: LM, TPP

Review & editing: LM, TPP

## Data and materials availability

All data are available in the main text or the supplementary materials.

## Code availability

No custom code beyond adaptation of existing software packages was used in this study.

All code will be provided upon reasonable request to the corresponding author.

---

**Extended Data Fig. 1.**
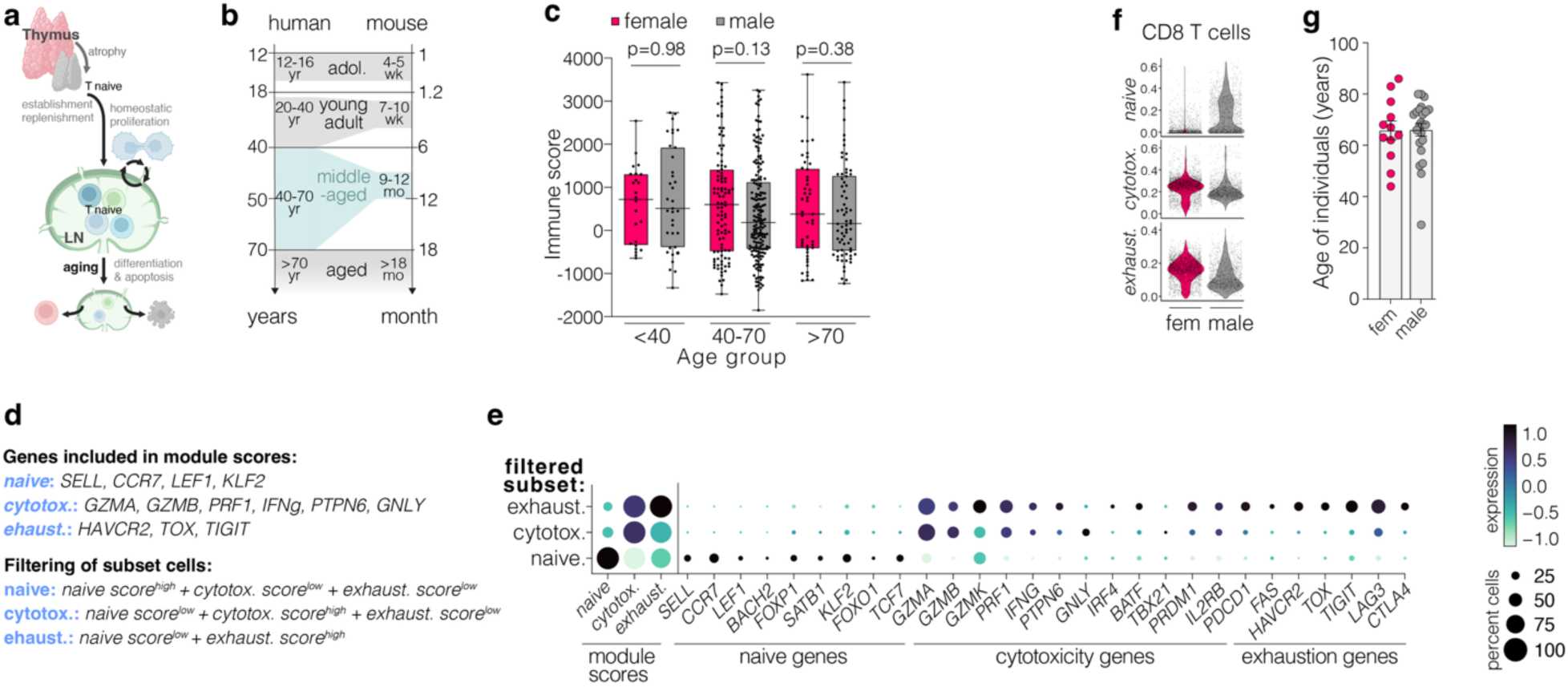
T cell infiltration in melanoma. **a,** schematic illustration of naive T cell and LN homeostasis. **b,** age categories in humans and mice^108,109^. **c,** immune scores derived from The Cancer Genome Atlas (TCGA) melanoma transcriptomes^110^, reflecting the estimated abundance of tumor-infiltrating immune cells. **d,** gene sets used to compute module scores for naive, cytotoxic, and exhausted CD8 T cell states. Integration of these scores was used to classify CD8 T cell subsets within tumor datasets. **e,** module score distributions and expression of canonical marker genes for each filtered CD8 T cell subset. Integrated data from Durante *et al.*^27^, Sade-Feldman *et al.*^28^, Li *et al.*^29^, Zhang *et al.*^30^. **f,** module score distributions for CD8 T cells from the Durante et al. dataset^27^. **g,** age distribution of melanoma patients included in the analyses shown in Fig. 1b-c (age provided when available; otherwise unspecified). Data shown as mean ± SEM with data points representing individual persons. Statistics calculated with MWU test (c).

**Extended Data Fig. 2.**
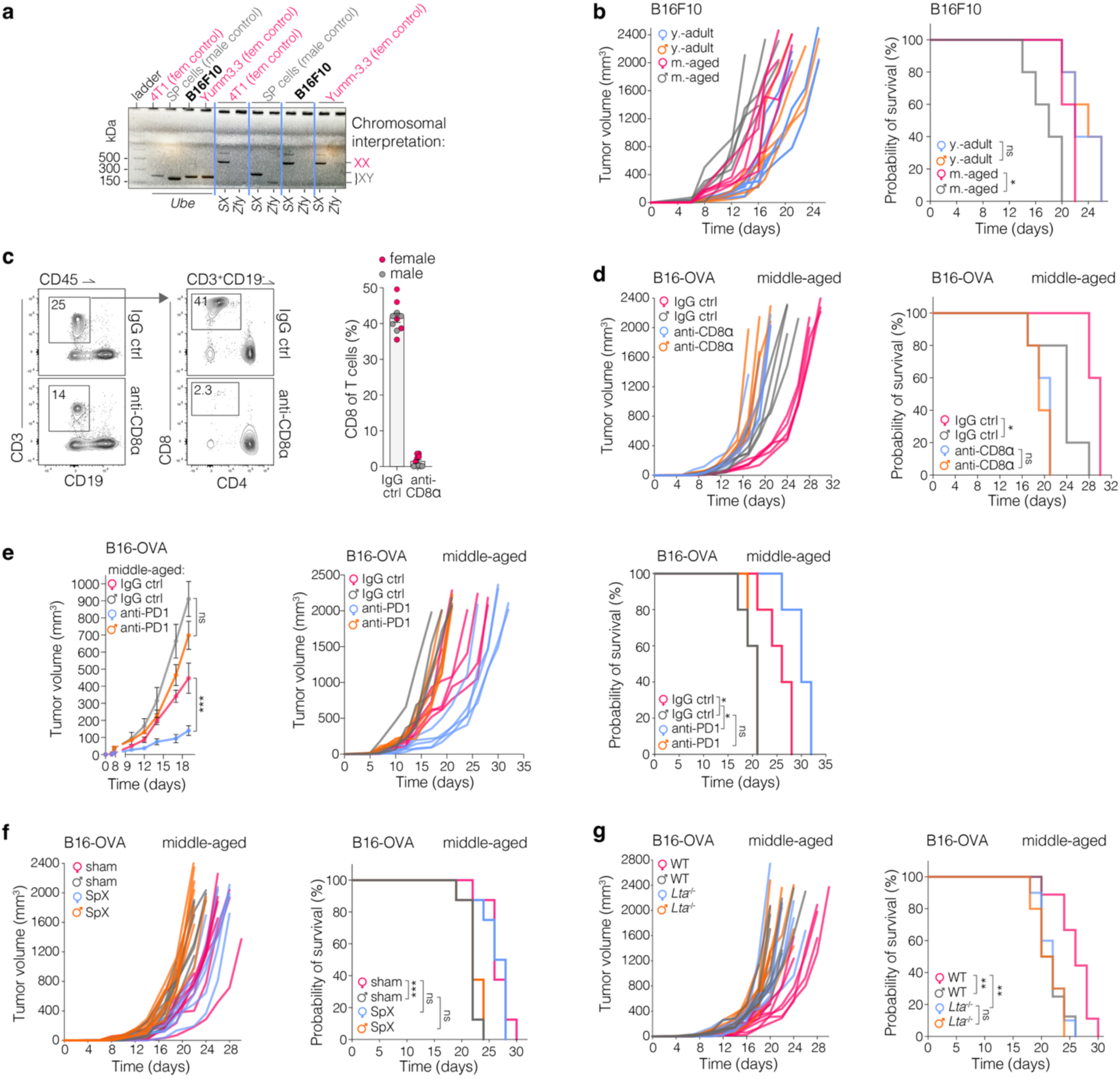
Melanoma tumor model growth analyses. **a,** PCR analysis, *Sly* intron 8 from the Y chromosome and *Xlr* intron 6 from the X chromosome (*SX*, 480/685 bp on XX chromosomes, 280 bp on Y chromosomes). Male control gene *Zfy* and Ubiquitin as general control gene *Uba1/Ube1y1* (Ube, 217 bp). The data show that B16F10 melanoma lack Y chromosome. **b,** B16F10 tumor growth and survival probability in adult and middle-aged mice (n=5-8/group), corresponding to ***left*** part of **Fig. 1d**. **c,** Flow cytometry of PBMCs from mice after treatment with IgG control or anti-CD8 antibody to deplete CD8 T cells, corresponding to **Extended Data Fig. 2d** and **Fig. 1d. d,** B16F10 tumor growth and survival probability in middle-aged mice after CD8 T cell depletion in comparison to controls. **e,** B16-OVA tumor growth and survival probability in middle-aged mice treated with IgG controls or anti-PD1 antibody (n=5/group). **f,** B16-OVA tumor growth and survival probability in middle-aged mice after sham surgery or splenectomy (SpX), n=8/group. **g,** B16-OVA tumor growth in middle-aged wild type (WT) or *Lta*^-/-^ mice (n=8-10/group). Data shown as mean ± SEM with data points representing individual animals. Statistics calculated with Log-rank test (b,d,e,f,g), 2-way ANOVA (e), ns=nonsignificant, *p<0.05,**p<0.01,***p<0.001.

**Extended Data Fig. 3.**
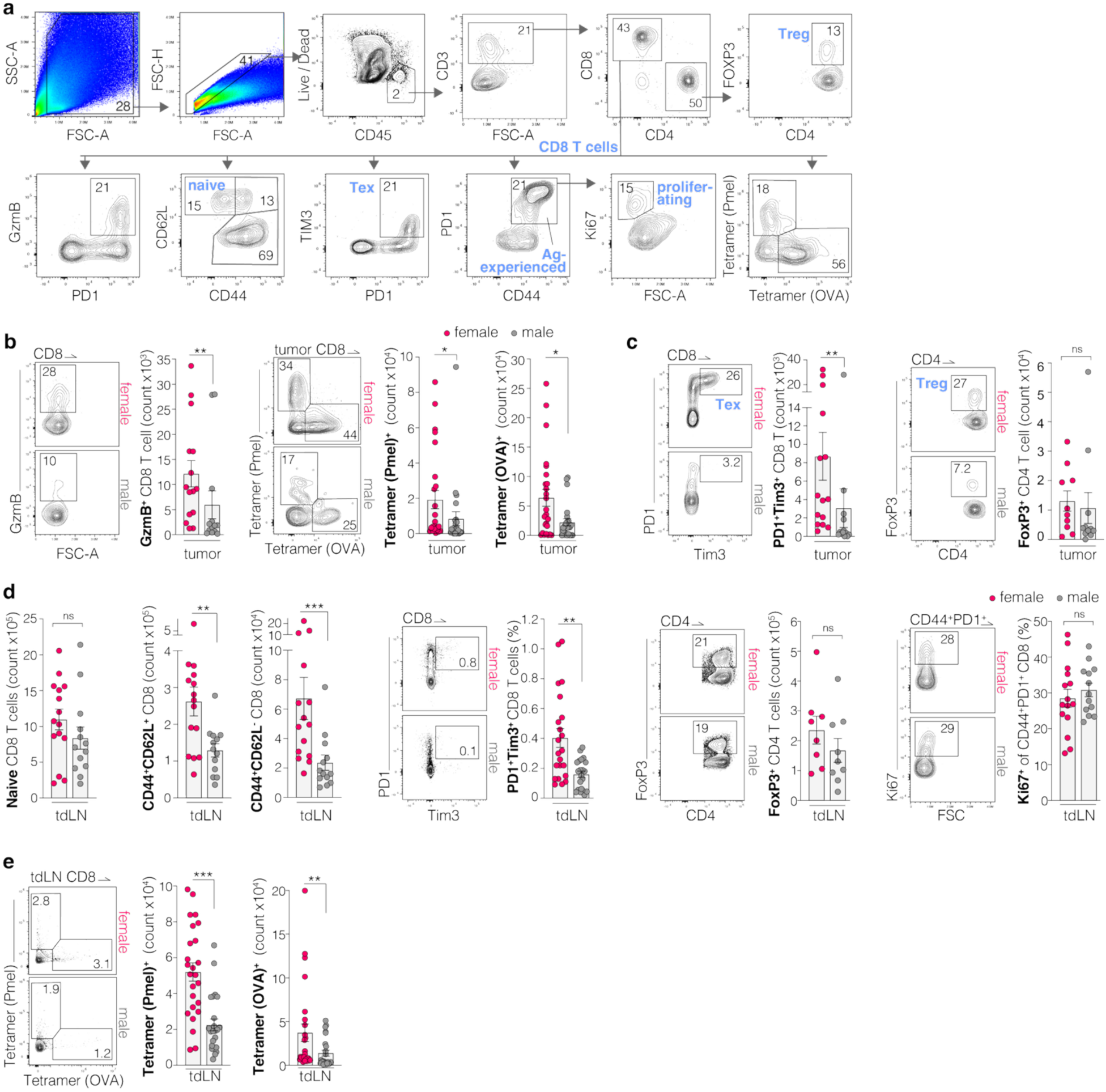
Flow cytometry analysis of the B16 mouse melanoma model. **a,** Flow cytometry gating strategy of lymphocytes in the primary tumor; regulatory T cells (Treg), exhausted T cells (Tex). **b and c** Flow cytometry analysis of CD8 T cells from the primary tumor of middle-aged mice implanted with B16-OVA tumors, group matched 13-15 days after implantation (corresponding to **Fig. 1e,f,g,i**)**. d,** Flow cytometry analysis of T cells in the tdLN of middle-aged mice implanted with B16-OVA tumors, group matched 13-15 days after implantation (corresponding to **Fig. 1e,f,g,i**)**. e,** Flow cytometry analysis of tetramer staining from melanoma draining LNs. Data points include the control sham/vehicle groups of middle-aged mice from thymectomy, gonadectomy and Degarelix treatment experiments **(Extended Data Fig. 12a, Extended Data Fig. 14c,g**). Data shown as mean ± SEM with data points representing individual animals. Statistics calculated with MWU test (b,c,d), ns=nonsignificant, **p<0.01, ***p<0.001.

**Extended Data Fig. 4.**
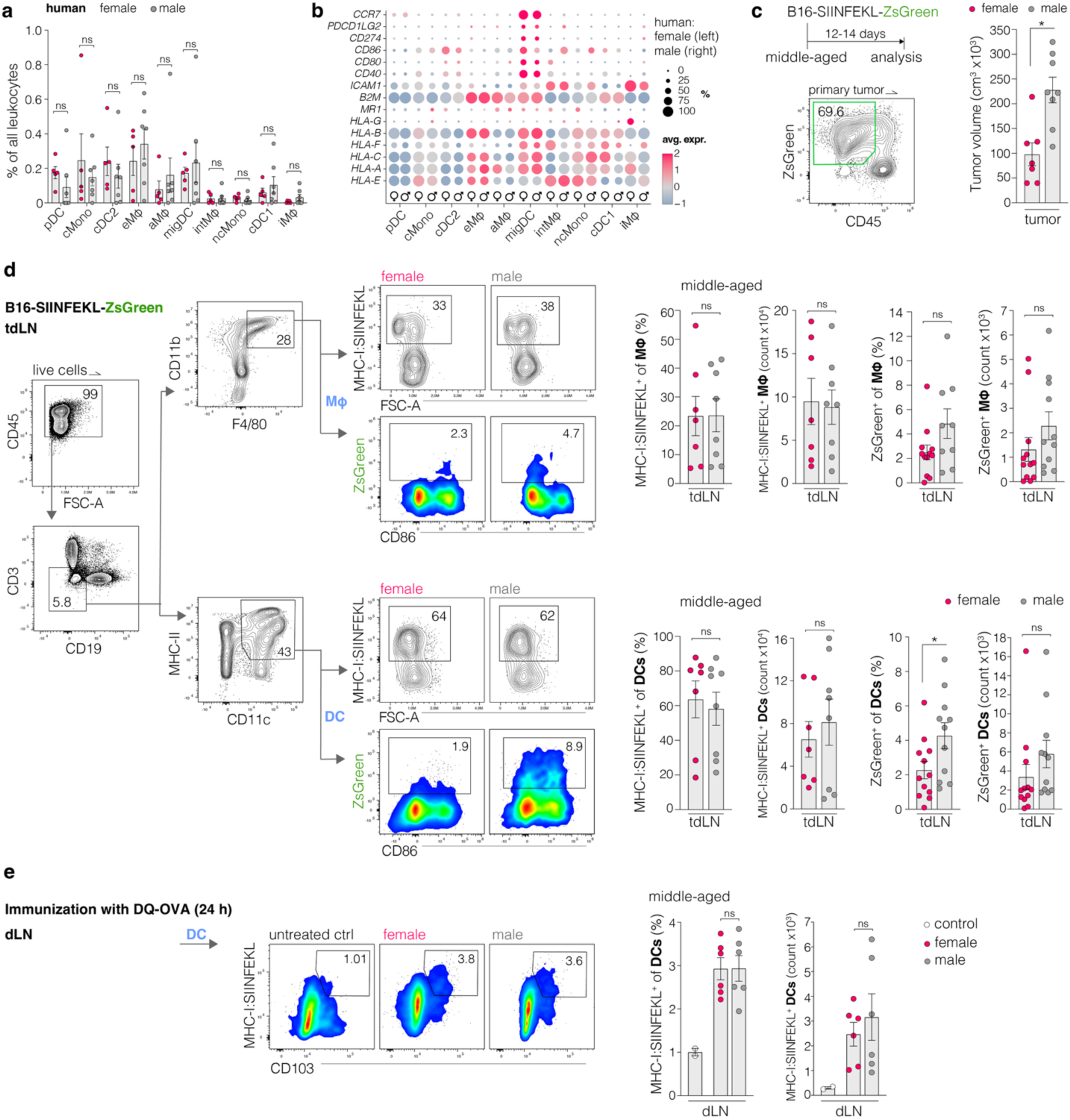
Subset distribution of APCs in human LNs and antigen presentation capacity in murine LNs. **a,** frequencies of subsets of APC in human LNs (53-73 years of age) taken from scRNA-seq data from Dominguez Conde *et al.*^48^ Plasmacytoid dendritic cell (pDC), classical monocytes (cMono), classical DC type 2 (cDC2), erythrophagocytic macrophage (eMΦ), alveolar (aMΦ), migratory DC (migDC), intestinal MΦ (intMΦ), non-classical Monocyte (ncMono), classical DC type 1 (cDC1), intermediate MΦ (iMΦ). **b,** expression levels of functional receptors in APCs of human LNs, females n=5, males n=7. **c,** B16-SIINFEKL-zsGreen tumors implanted in middle-aged mice, volume at day of analysis and representative flow cytometry plot demonstrating presence of zsGreen in tumor cells and in some CD45^+^ leukocytes within the primary tumor. **d,** representative plots of the flow cytometry analysis and quantification of DCs and MΦ presenting SIINFEKL peptide via MHC-I (MHC-I:SIINFEKL) or carrying tumor antigen (zsGreen) in tdLNs of middle-aged mice with B16-SIINFEKL-zsGreen tumors. **e,** representative plots of the flow cytometry analysis and quantification of MHC-I:SIINFEKL^+^ in draining LNs (dLN) of middle-aged mice 25 hours after immunization with DQ-OVA, which identifies antigen uptake and processing. Data shown as mean ± SEM with data points representing individual animals or persons. Statistics calculated with MWU test (a,c,d,e), ns=nonsignificant, *p<0.05.

**Extended Data Fig. 5.**
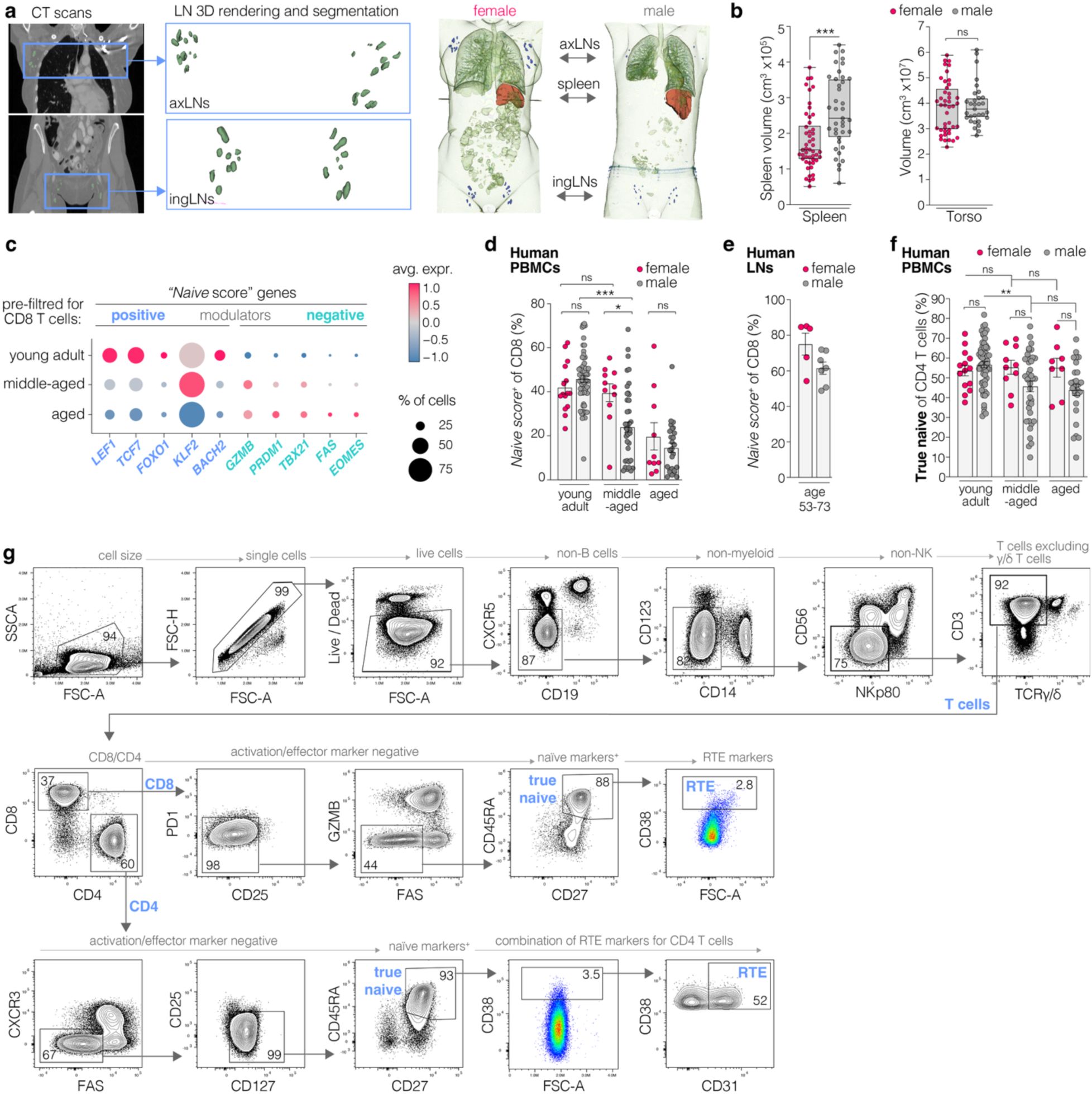
Human LN volumes and naive CD8 T cells. **a,** Schematic representation of LN segmentation, 3D rendering, and quantification of LN volumes based on CT scans from Roth et al. and Rister et al.^38,39^. **b,** Data corresponding to **Fig. 2a,b**, Segmented and quantified spleen and torso volumes. **c,** Expression levels of genes included in the “*Naive Score*” with genes that positively or negatively modulate the score. **d,** Quantification of naive CD8 T cells in human PBMCs using the *Naive Score*, based on scRNA-seq data from Terekhova *et al.*^46^. **e,** Quantification of naive CD8 T cells in human LNs using the *Naive Score*, based on scRNA-seq data from Dominguez Conde *et al.* ^48^. **f,** Flow cytometry quantification of true naive CD4 T cells from human PBMCs, public data set^47^. Gating Strategy in **Extended Data Fig. 5g**. **g,** Flow cytometry gating strategy for human PBMCs, including recent thymic emigrants (RTEs), with data derived from Bohacova & Terekhova *et al.*^47^. Data shown as mean ± SEM with data points representing individual animals or human subjects. Statistics calculated with MWU test (b,d,f), ns=nonsignificant, *p<0.05, **p<0.01, ***p<0.001.

**Extended Data Fig. 6.**
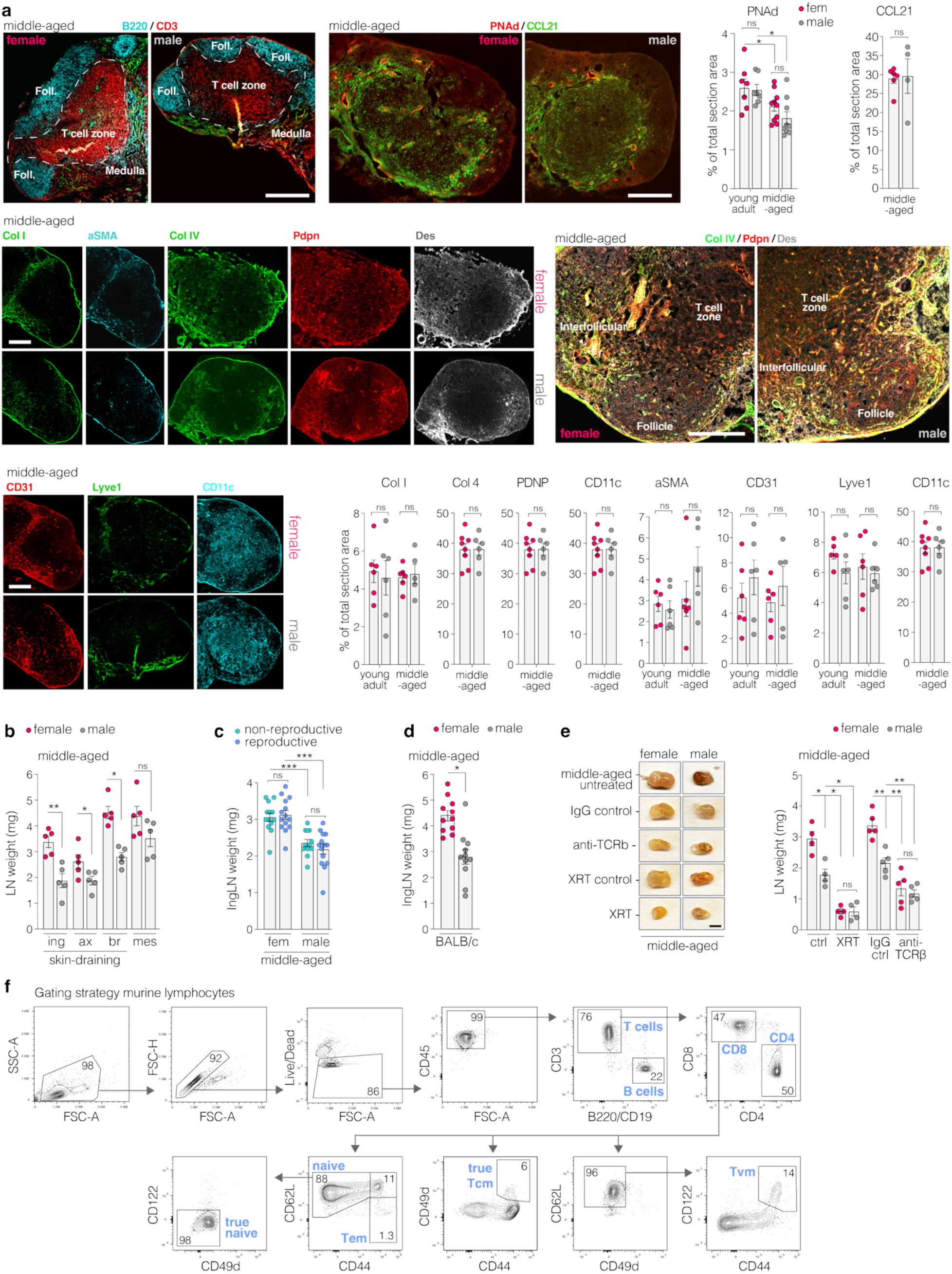
Age-and sex-related contraction of LNs with and without lymphodepletion. **a,** Representative images and quantification of inguinal LN sections; B220 (B cells), CD3 (T cells), peripheral node adressin (PNAd, high endothelial venules), C-C motif chemokine ligand 21 (CCL21), collagen I (Col I), alpha smooth muscle actin (αSMA), Col IV, podoplanin (Pdpn), Desmin (Des), CD31 (PECAM-1, endothelial cells), lymphatic vessel endothelial hyaluronan receptor-1 (Lyve-1, lymphatic endothelial cells), CD11c (dendritic cells). Scale bar: 200 µm, 50 µm (FRC network images). **b,** LN weights in different age groups of female and male mice, including inguinal (ing), axillary (ax), brachial (br), and mesenteric (mes) LNs. **c,** Weights of inguinal LNs in non-reproductive and reproductive (retired breeder) female and male mice. **d,** LN weights in middle-aged female and male BALB/c mice. **e,** Representative photographs and weight quantification of inguinal LNs from middle-aged mice treated with irradiation (XRT, sublethal dose of 4 Gy) or anti-TCRβ antibody-mediated T cell depletion. Scale bar 200 µm and 50 µm (FRC network images). **f,** Flow cytometry gating strategy for murine lymphocytes, including central memory T cells (Tcm), effector memory T cells (Tem), and virtual memory T cells (Tvm). Data shown as mean ± SEM with data points representing single LNs of individual animals. Statistics calculated with MWU test (a,b,c,d,e), ns=nonsignificant, *p<0.05, **p<0.01, ***p<0.001.

**Extended Data Fig. 7.**
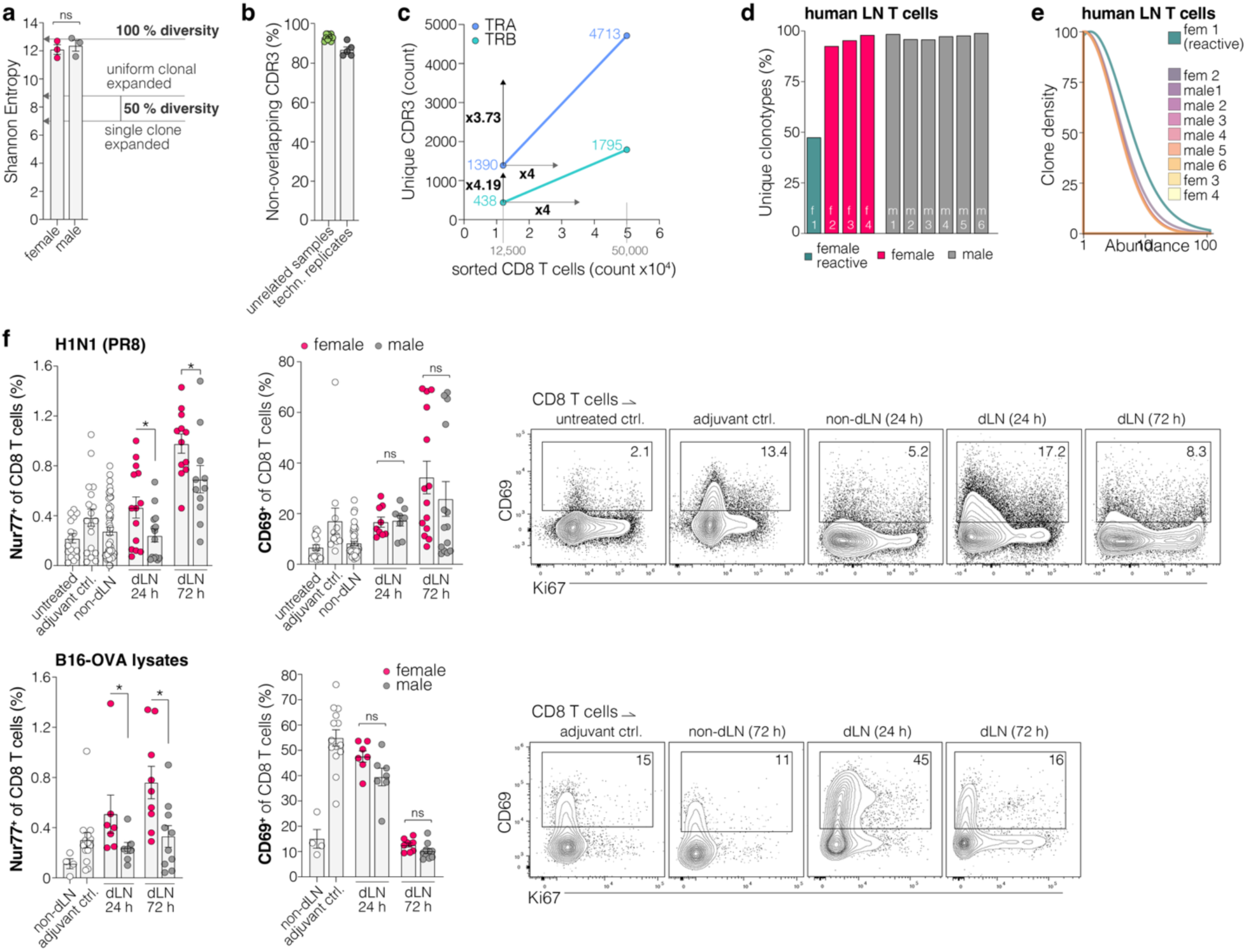
TCR diversity of naive T cells in LNs and T cell activation after immunization in mice. **a,** Shannon Entropy analysis of the TCR repertoire (T cell receptor beta chain, TRB) diversity using bulk TCR sequencing data of CD8 T cells sorted from inguinal LNs of middle-aged mice. Indicated are approximations of theoretical values with 100% diversity and 50% diversity in reactive LNs with an even clonal expansion or a single expanded clone. **b,** CDR3 (TRB) sequences overlapping with other samples or technical replicates of the same sample. **c,** Bulk TCR sequencing of T cells from the same LN sample (ingLN of middle-aged male) with different CD8 T cell counts (12,500 and 50,000 cells). Indicated are unique CDR3 (TRA, TRB) and the fold change (x…) between samples with different input cell counts. Results show that almost every additional cell in this LN sample equals an additional unique TCR. **d,** ScTCR-seq of T cells from human LNs, data set from Dominguez Conde *et al.*^48^ Percent of unique clonotypes (including combination of TRA, TRB). One female sample with apparent reactive LN (see **e**). **e,** Clonal density and abundance (arbitrary units) of samples depicted in **d**. **f,** Flow cytometry analysis of Nur77 and CD69 expression in CD8 T cells 24 or 72 hours after immunization with inactivated H1N1 (PR8) virus particles or B16-OVA tumor cell lysates. Data shown as mean ± SEM with data points representing single LNs of individual animals or persons. Statistics calculated with t test (a), MWU test (f), ns=nonsignificant, *p<0.05, ***p<0.001.

**Extended Data Fig. 8.**
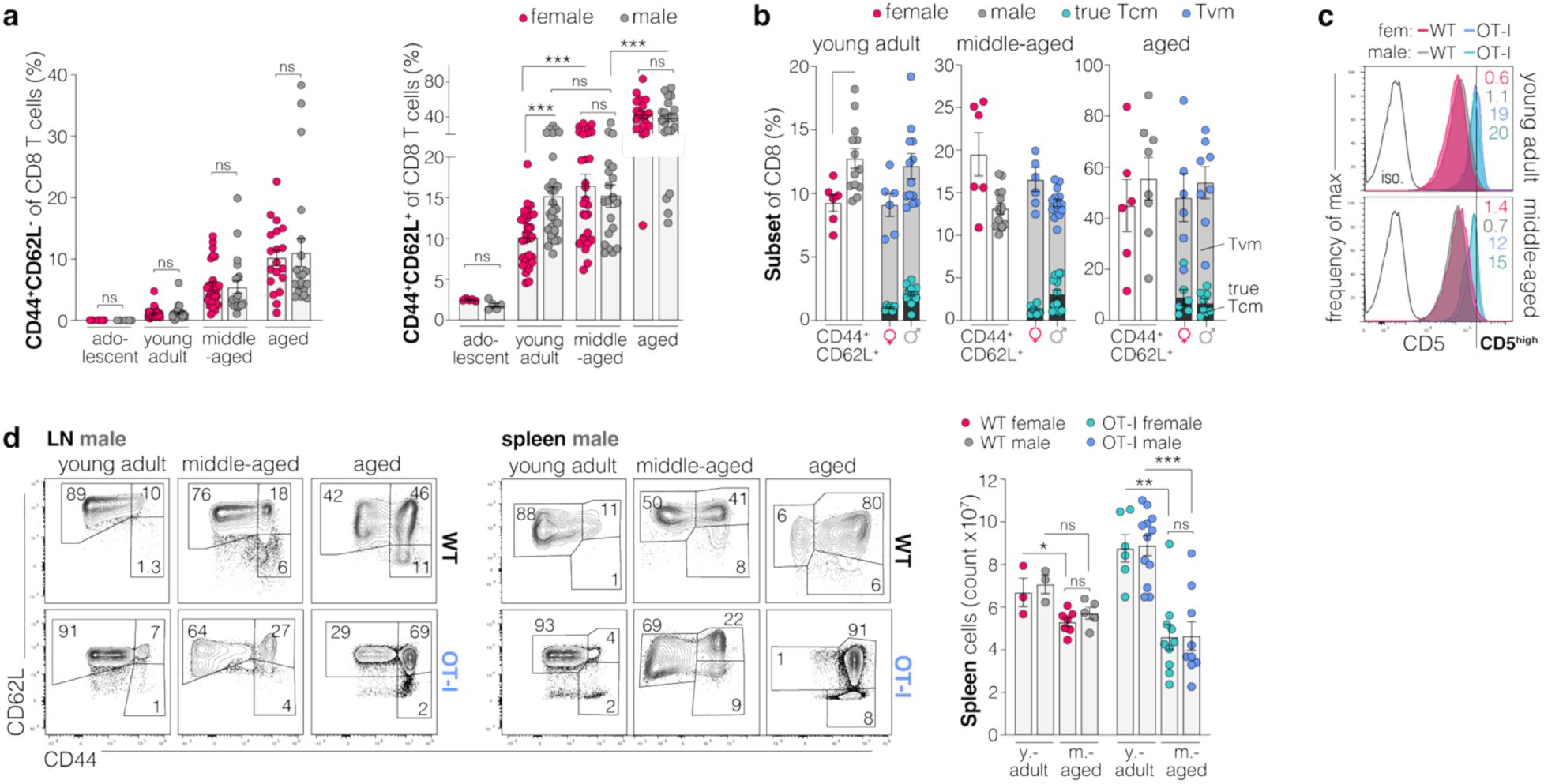
Age group related analysis of naive and memory CD8 T cells in wild type and OT-I mice. **a,** Flow cytometry analysis of central memory-like (CD44^+^CD62L^+^) and effector-like (CD44^+^CD62L^-^) CD8 T cells in LNs of wild-type (WT) mice. **b,** Quantification of flow cytometry analysis of CD8 T cells from the ingLN of adult, middle-aged and aged mice show that CD44^+^CD62L^+^ contain both true Tcm and virtual memory (Tvm) cells, with a majority being Tvm. **c,** Histograms of CD5 expression in true naive CD8 T cells, vertical line indicates CD5^high^ cells from ingLNs of WT or OT-I mice, young adults (y. adult) and middle-aged (m.-aged). **d,** Representative flow cytometry plots of CD8 T cells and quantification of total cell count from the spleens of WT and OT-I mice. Data shown as mean ± SEM with data points representing single LNs or tissues of individual animals. Statistics calculated with MWU test (a,d), ns=nonsignificant, *p<0.05, **p<0.01, ***p<0.001.

**Extended Data Fig. 9.**
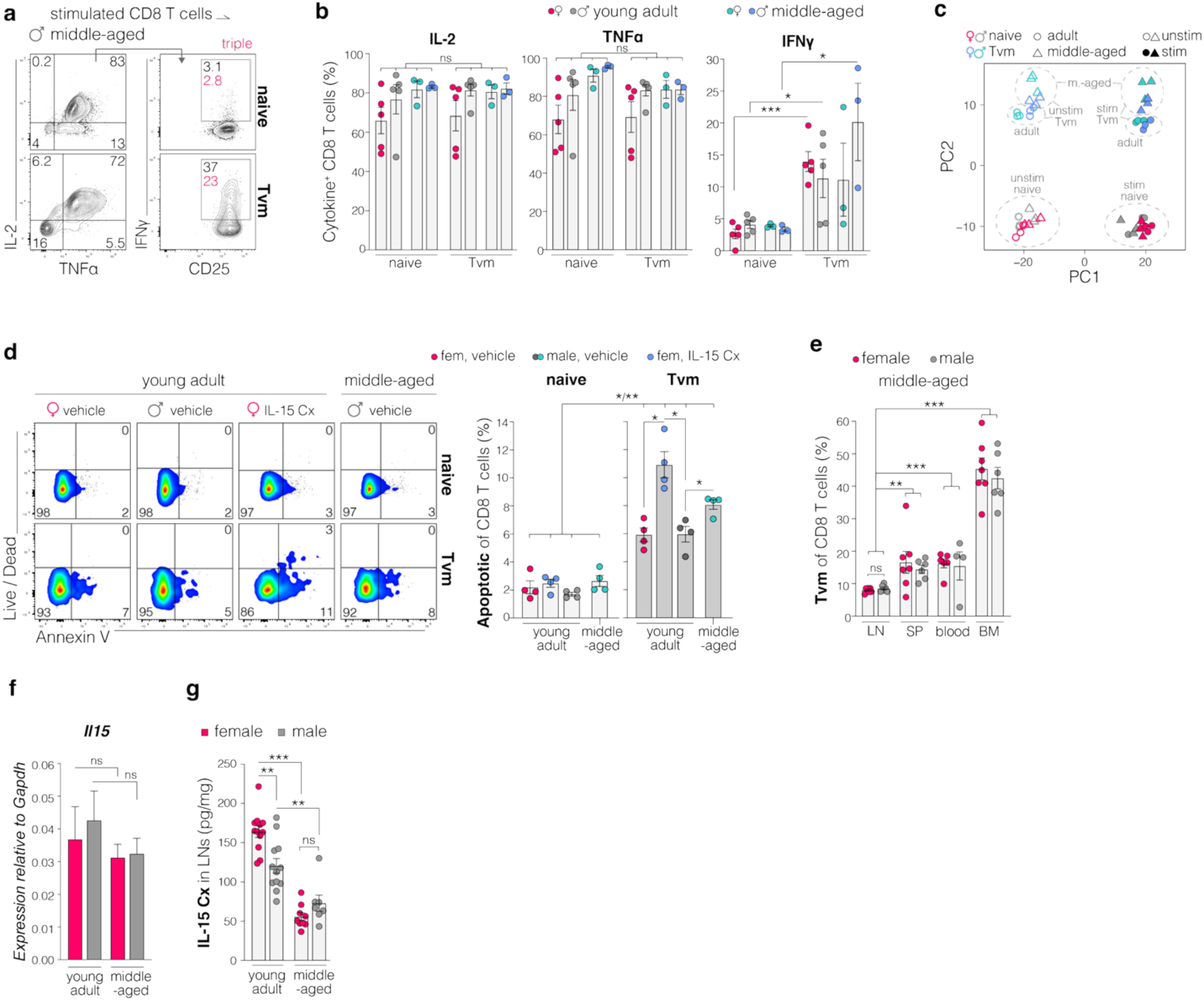
CD8 Tvm cells in young-adult, middle-aged and aged mice. **a,** Example plots and b, quantification of cytokine production in subsets of CD8 T cells after in vitro following stimulation. **c**, Principal component (PC) analysis of VST-normalized bulk RNA-seq counts across all 48 samples. **d,** Flow cytometry analysis of Live/Dead (Zombie UV) and Annexin-V staining in true naive or Tvm cells of young adult mice after treatment with vehicle solution or IL-15 Cx or from vehicle treated middle-aged males. Analysis 1 week after end of treatment. **e,** Flow cytometry analysis of Tvm CD8 T cells in LNs, spleen (SP), blood and bone marrow (BM). **f,** qPCR gene expression analysis of *Il15* relative to *Gapdh* in murine ingLN, n=10-15/group. **g,** ELISA quantification of IL-15/IL-15 receptor alpha complex (Il-15 Cx) in ingLNs of young adult and middle-aged mice, indicated relative to weight of LNs. Data shown as mean ± SEM with data points representing single LNs or tissues of individual animals. Statistics calculated with Student’s t-test (b) or MWU test (d,e,f,g), ns=nonsignificant, *p<0.05, **p<0.01, ***p<0.001.

**Extended Data Fig. 10.**
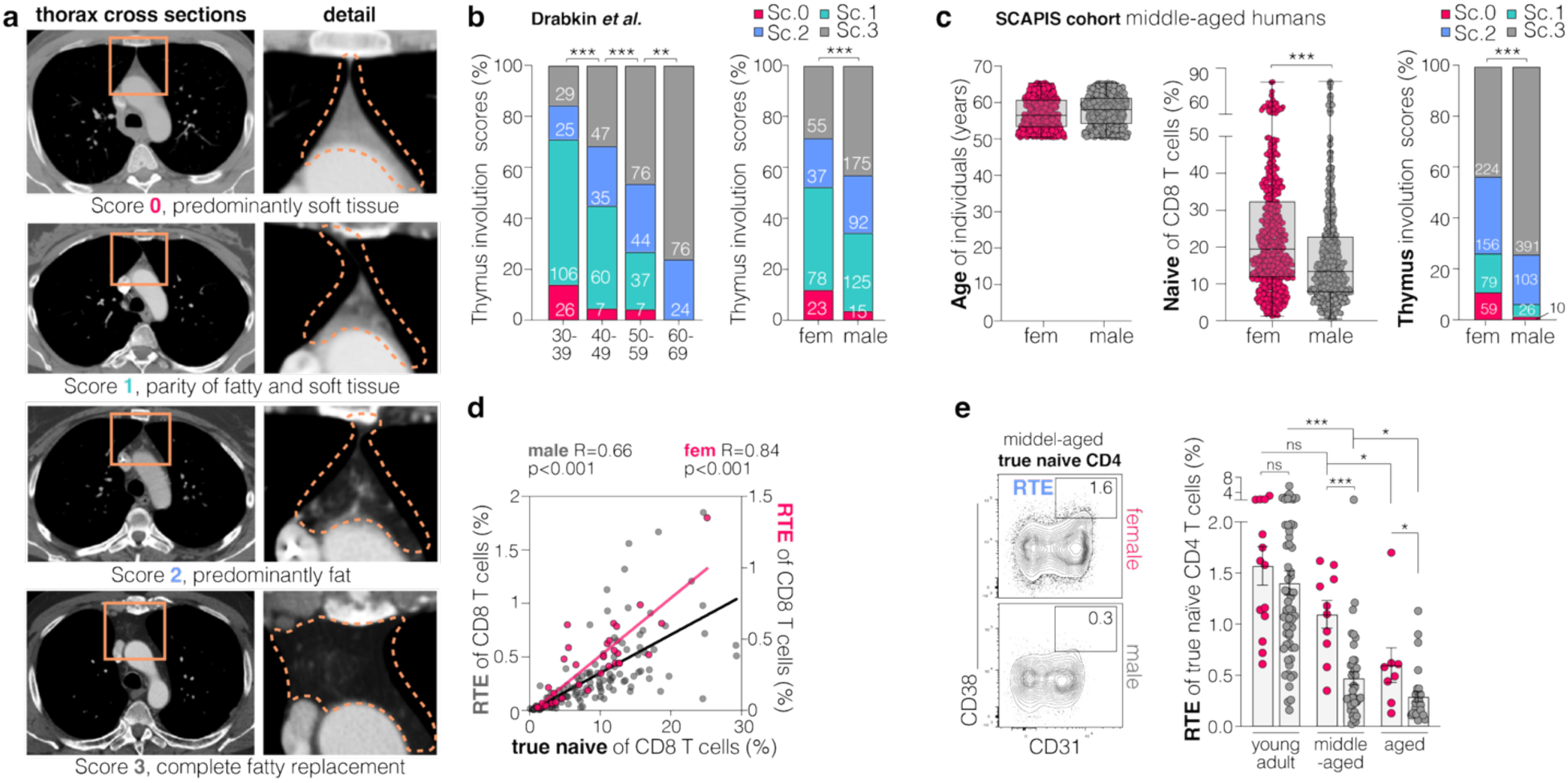
Human thymus and circulating recent thymic emigrants. **a,** representative images of CT scans and scores to evaluate the involution of human thymuses, score 0: predominantly soft tissue; score 1: parity of fatty and soft tissue; score 2: predominantly fat; score 3: complete fatty replacement. **b,** data published by Drabkin *et al.*^72^ confirming age-related thymus involution using the scoring system as described in **a**. Numbers in columns indicate number of individuals per group. **c,** age in years (all middle-aged, 50-65 years old), proportion of naive CD8 T cells within the CD8 T cell compartment in blood samples and thymus involution scores of the SCAPIS cohort of middle-aged individuals, n=530 females and 518 males. **d,** correlation of the frequency of the true naive subset and of RTE relative to CD8 T cells in public data set^47^. **e,** representative flow cytometry plot and quantification of CD4 RTE and true naive CD4 T cells in human blood, public data set^47^. Gating strategy in **Extended Data Fig. 5g**. Data shown as mean ± SEM with data points representing individual human subjects. Statistics calculated with Fisher’s exact test (b,c), Pearson correlation (d), MWU test (c,e), ns=nonsignificant, *p<0.05, **p<0.01, ***p<0.001.

**Extended Data Fig. 11.**
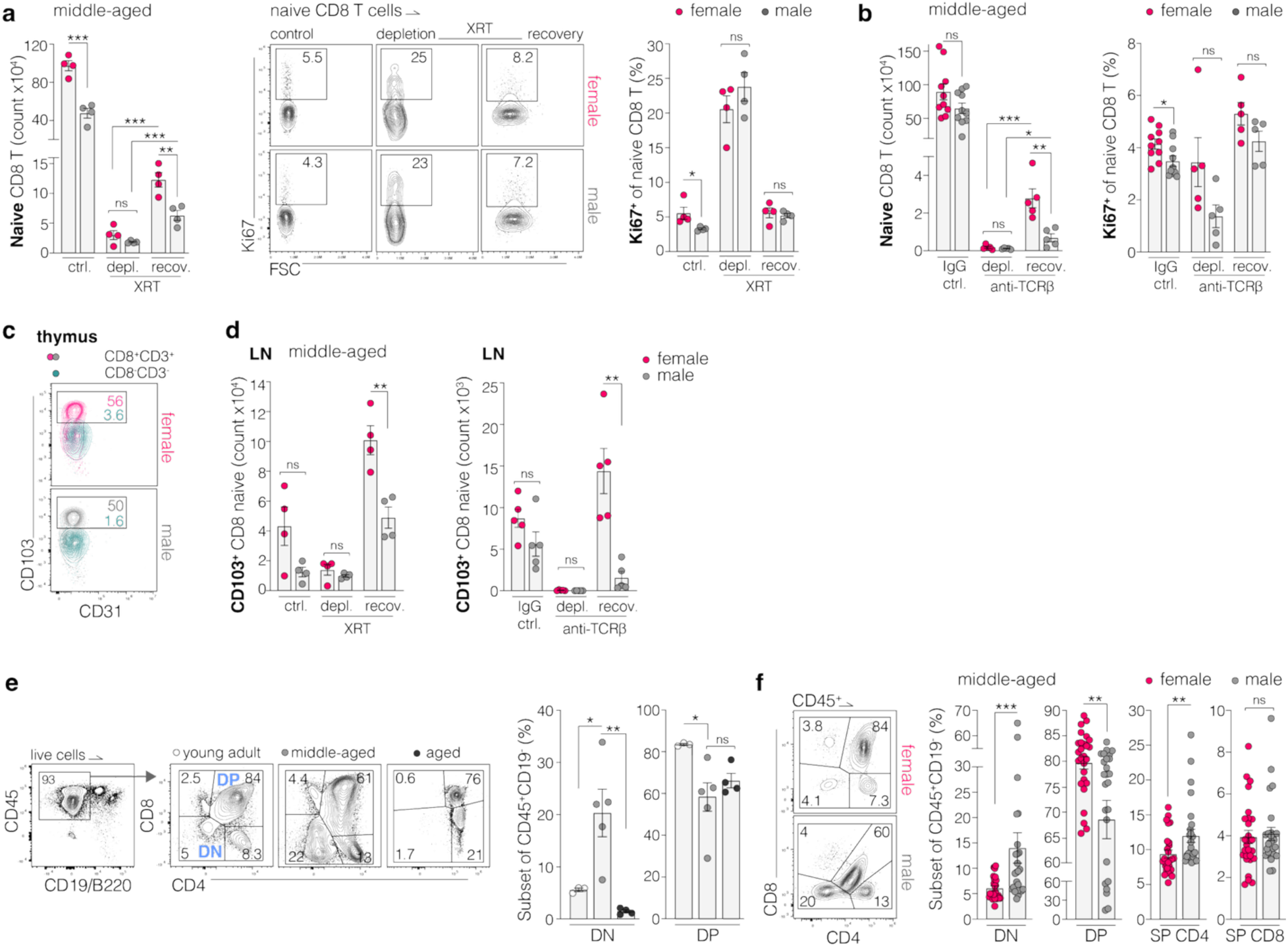
T cell recovery after experimental lymphodepletion and thymocyte alterations during aging. **a,** Flow cytometry analysis of naive CD8 T cells and proliferation of naive T cells (Ki67^+^) in middle-aged untreated control mice or after irradiation (XRT, sublethal dose with 4 Gy) induced lympho-depletion (depl., day 7) and after a period of recovery (recov., day 28). **b,** Naive CD8 T cell counts and Ki67 expression in middle-aged mice treated with IgG control or anti–TCRβ antibody. Shown are depletion at day 7 and recovery at day 19. **c,** Representative flow cytometry plots of CD103 expression in CD8⁺CD3⁺ single-positive (SP) thymocytes, marking recent thymic emigrant (RTE) precursors. CD8⁻CD3⁻ cells serve as gating controls. **d,** Frequencies and absolute numbers of CD103⁺ naive CD8 T cells in inguinal LNs of middle-aged mice at depletion (day 7) and recovery phases after XRT or anti–TCRβ treatment. **e,** Flow cytometry analysis of thymocytes (CD45^+^CD19^-^/B220^-^) in male mice of different age groups; including double negative (DN) and double positive (DP) cells. **f,** flow cytometry analysis of thymocytes (CD45^+^CD19^-^) in middle-aged female and male mice; including DN and DP cells and single positive (SP) cells. Data shown as mean ± SEM with data points representing individual mice. Data shown as mean ± SEM with data points representing single LNs or thymuses of individual animals. Statistics calculated with MWU test (a,b,d,e,f), ns=nonsignificant, *p<0.05, **p<0.01, ***p<0.001.

**Extended Data Fig. 12.**
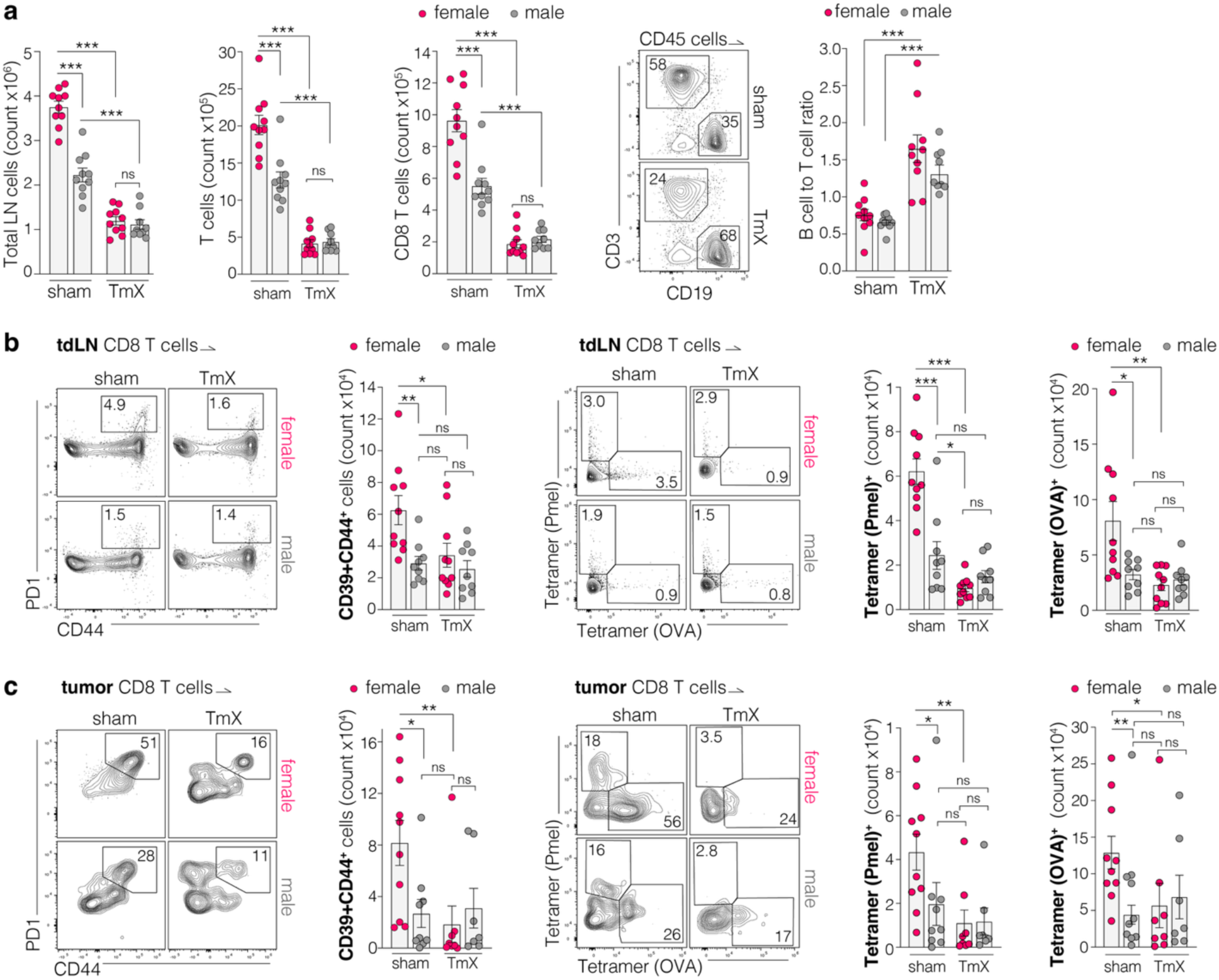
Consequences of thymectomy for LNs and tumor recognition in tdLN and within the primary tumor. **a,** quantification of total LN cells, T cells, CD8 T cells and the T cell to B cell ratio in ingLNs of middle-aged sham and TmX treated mice. **b,** flow cytometry analysis of CD8 T cells in tdLNs of sham and TmX mice with B16-OVA tumors. Corresponding to **Fig. 5i,j,k**. **c,** flow cytometry analysis of CD8 T cells from the primary B16-OVA tumor of sham and TmX mice. Corresponding to **Fig. 5i,j,k**. Data shown as mean ± SEM with data points representing single LNs or tumors of individual animals. Statistics calculated with MWU test (a,b,c), ns=nonsignificant, *p<0.05, **p<0.01, ***p<0.001.

**Extended Data Fig. 13.**
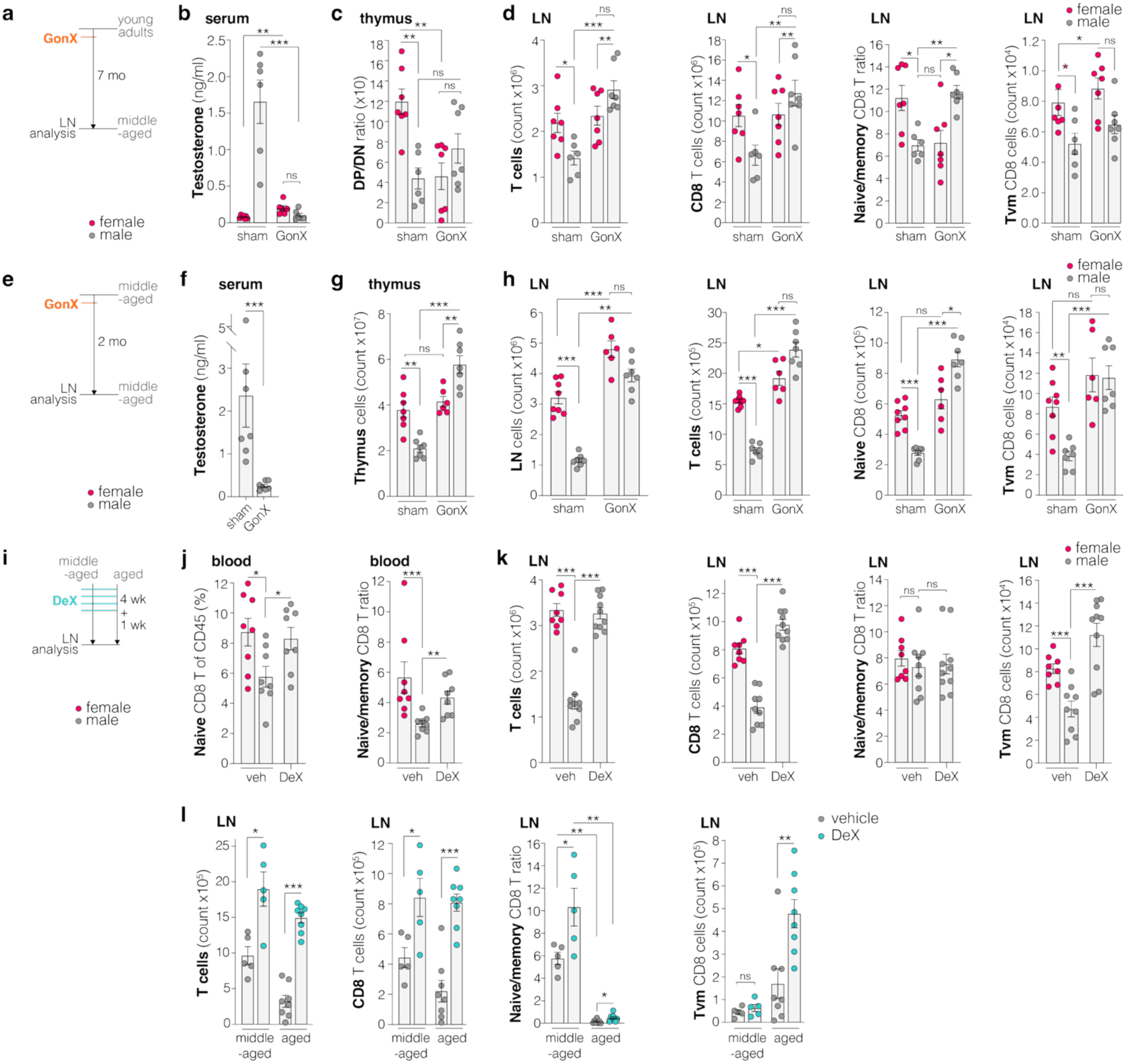
Consequences of sex steroid ablation for the thymus and lymph nodes. **a,** Experimental setup, Gonadectomy (GonX) in adult mice and analysis middle-aged. **b,** Analysis of serum testosterone levels. **c,** Flow cytometry analysis of thymocytes (CD45^+^CD19^-^/B220^-^); ratio of double positive (DP) and double negative (DN) cells. **d,** Flow cytometry analysis and quantification of T cells from tumor-naive LNs. **e,** Experimental setup, GonX in middle-aged mice and analysis two months after surgery. **f,** Analysis of serum testosterone levels. **g,** Number of total thymocytes. **h,** LN cellularity and flow cytometry analysis of T cells from tumor-naive LNs. **i,** Experimental setup, middle-aged and aged mice were treated with vehicle (veh) or Degarelix (DeX). **j,** Flow cytometry analysis of naive and memory CD8 T cells in blood of middle-aged mice. **k,** Flow cytometry analysis of T cells in tumor-naive LNs from middle-aged mice. **l,** Flow cytometry analysis of T cells from tumor-naive LNs in male mice. Data shown as mean ± SEM with data points representing single LNs, blood samples or thymuses of individual animals. Statistics calculated with MWU test (b,c,d,f,g,h,j,k,l), ns=nonsignificant, *p<0.05, **p< 0.01, ***p<0.001.

**Extended Data Fig. 14.**
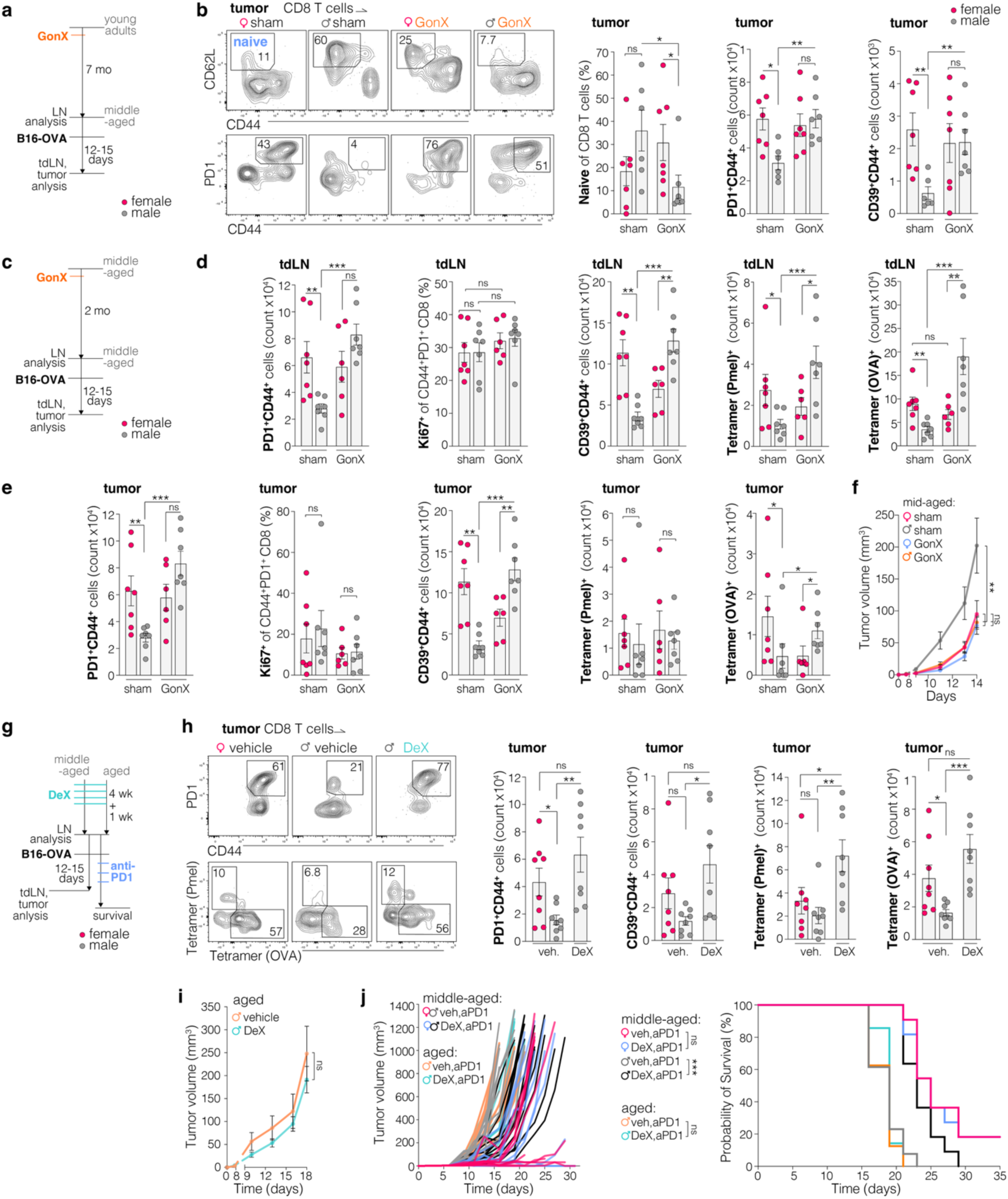
Consequences of sex steroid ablation tumor recognition in tdLNs and the primary tumor. **a,** Experimental setup, gonadectomy (GonX) in young adults followed by B16-OVA tumor implantation and analysis in middle-aged mice. **b,** Flow cytometry analysis of T cells in tumors. **c,** Experimental setup, GonX in middle-aged mice and analysis or B16-OVA tumor implantation two months after surgery. **d,** Flow cytometry analysis of tdLNs and **e,** TILs. **f,** B16-OVA tumor growth in mice after sham or GonX surgery, n=7/group. **g,** Experimental setup, left, young adult and middle-aged mice were gonadectomized (GonX, ovariectomy for females, orchiectomy in males) or, right, middle-aged and aged mice were treated with vehicle (veh) or Degarelix (DeX) before analysis of tumor-naive LNs, or implantation of B16-OVA tumors with anti-PD1 antibody treatment. **h,** Flow cytometry analysis of TILs. **i,** B16-OVA tumor growth in vehicle or DeX treated aged mice (n=4-5/group). **j,** B16-OVA tumor growth and survival probability in middle-aged and aged mice after treatment with DeX and anti-PD1 (n=7-13/group). Data shown as mean ± SEM with data points representing single LNs or tumor of individual animals. Statistics calculated with MWU test (b,g,e,h), two-way repeated measures ANOVA (f,i), or Log-rank test (j) ns=nonsignificant, *p<0.05, **p<0.01, ***p<0.001.

**Extended Data table 1.**
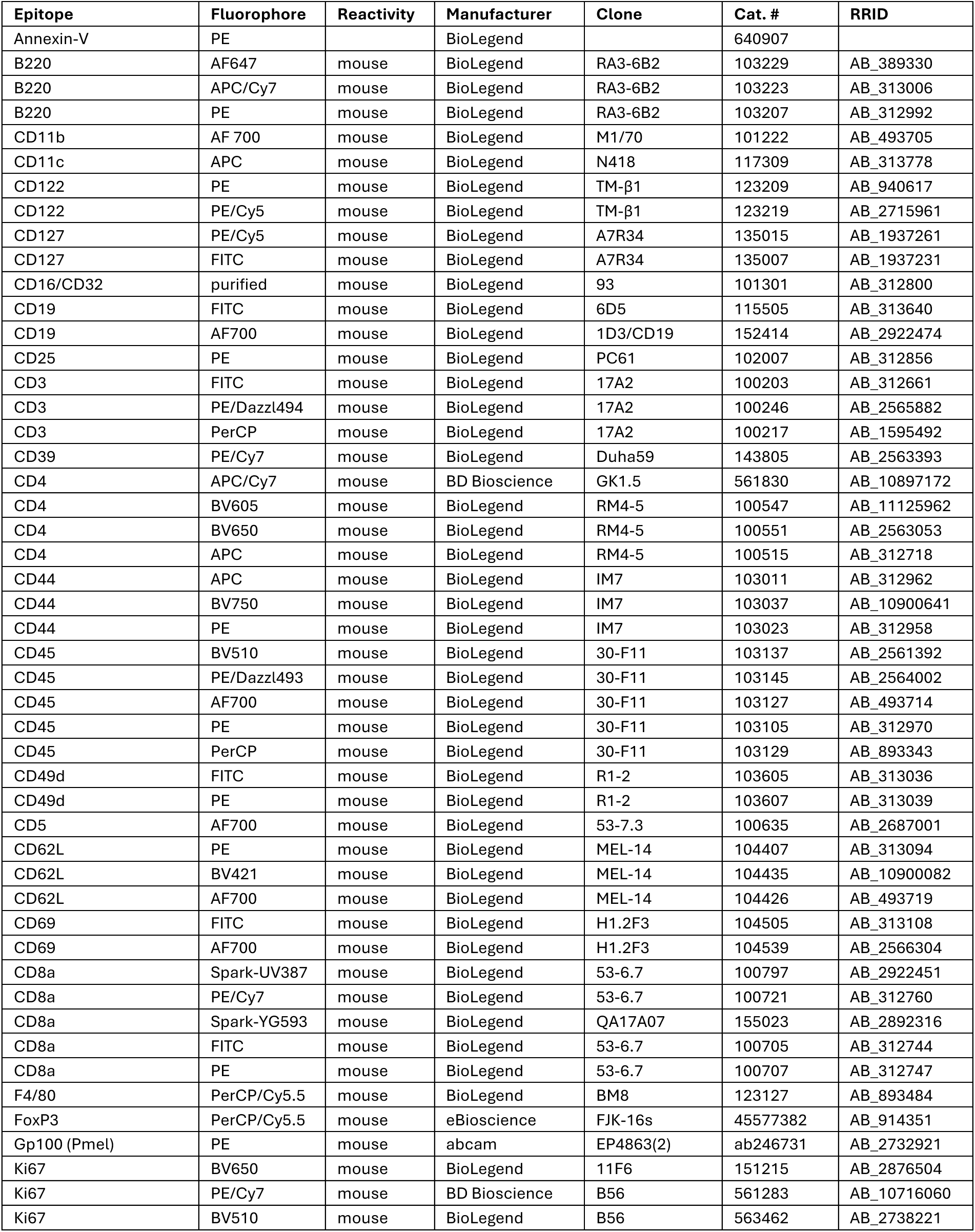
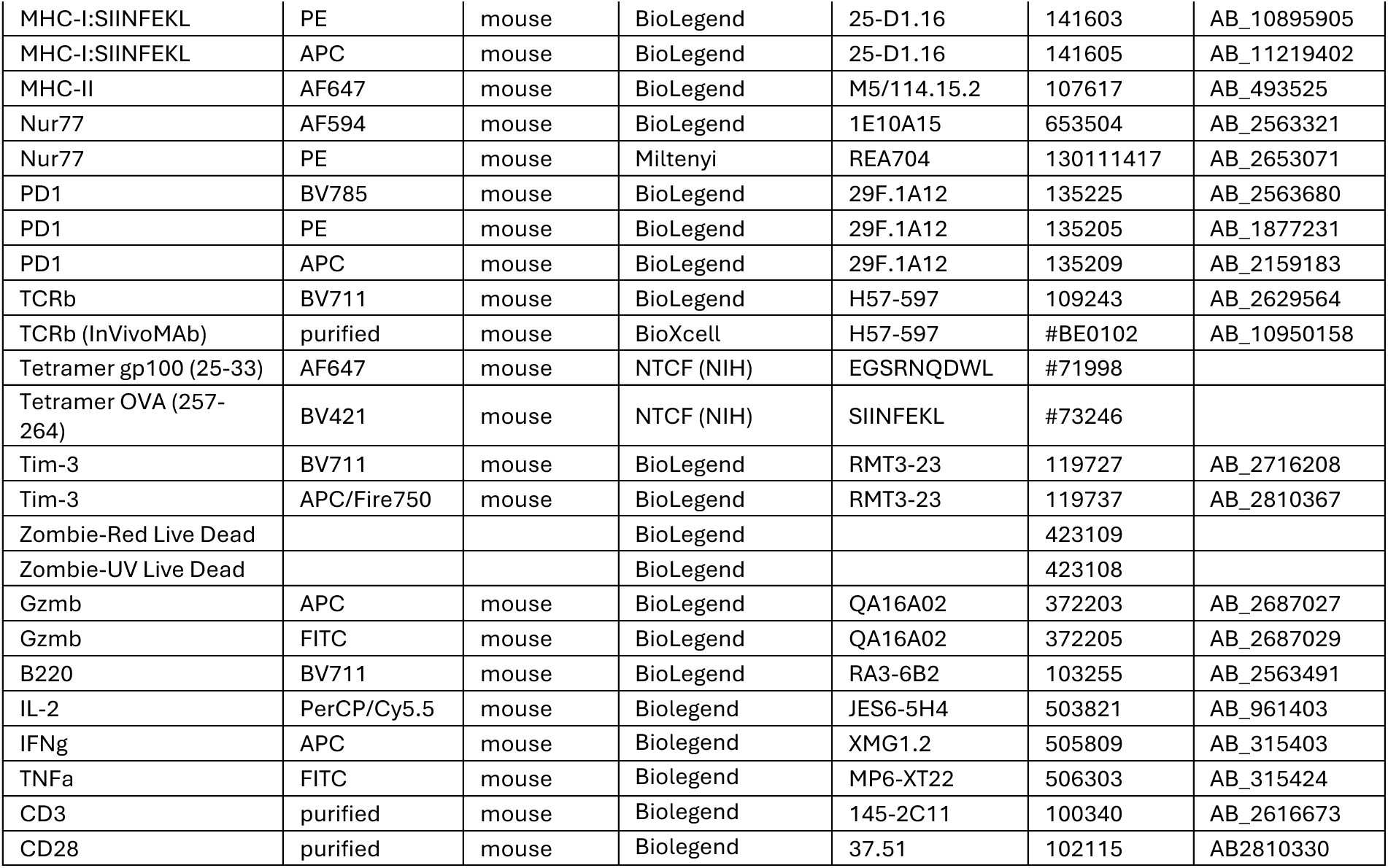
Antibodies, tetramers, dyes.

**Extended Data table 2.**
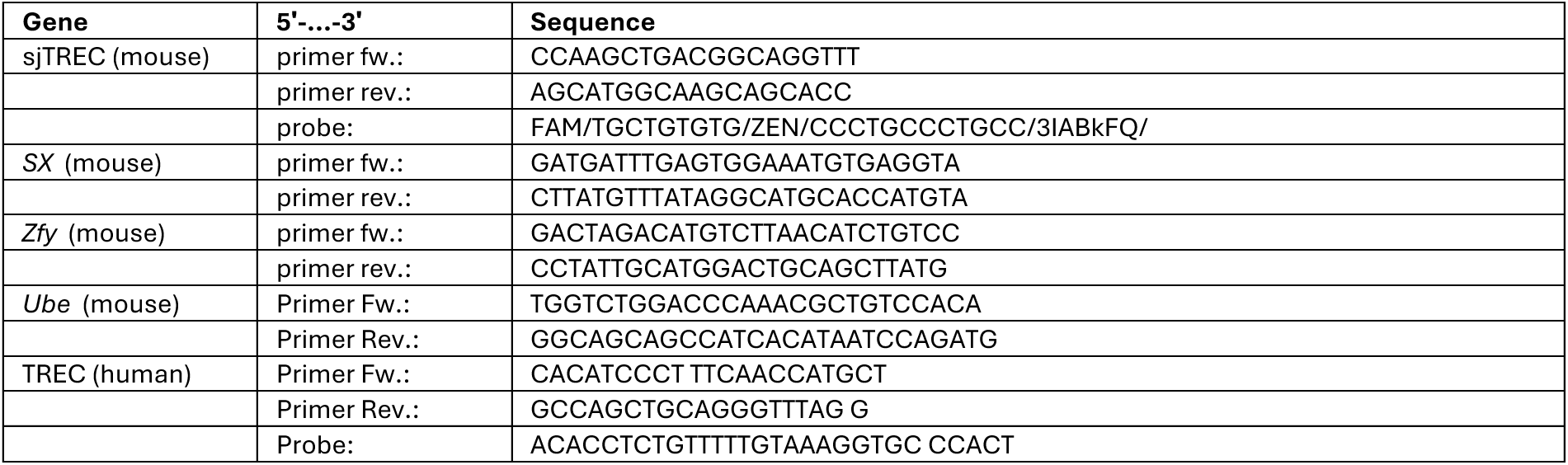
Primers and probes. Abbreviations: sjTREC (signal joint T cell receptor excision circle), Fw. (forward), rev. (revers).

