## Supplementary Informations for "Lymph node contraction links sex-biased naive CD8 T cell decline to compromised antigen recognition during middle age"

### Supplementary Information

**a**

$t$ = *time step* (1 time step = 3 weeks = month x 4.345/3)
$N_t$ = fraction of naive cells remaining at time $t$
$k_{sex}$ = conversion probability per step (sex specfic)
$C_t$ = fraction converted into Tvm
$s_{sex}(t)$ = thymic input at time t

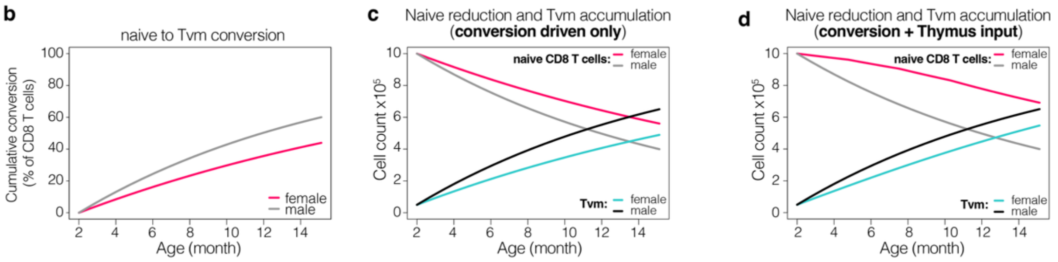


**Core assumptions:**
- fixed fraction ($k$) of naive converts per time step
- no replenishment of naive T cells
- $V_0$ = 5x10^5 Tvm cells
- $k_{female}$ = 0.03
- $k_{male}$ = 0.03 × (1.1 / 0.7) = 0.047

**Core assumptions:**
- fixed fraction ($k$) of naive converts per time step
- no replenishment of naive T cells
- Tvm conversion 1:1 naive reduction
- $N_0$ = 10^6 naive cells
- $V_0$ = 5x10^5 Tvm cells
- $k_{female}$ = 0.03
- $k_{male}$ = 0.03 × (1.1 / 0.7) = 0.047

**Core assumptions:**
- fixed fraction ($k$) of naive converts per time step
- Tvm conversion 1:1 naive reduction
- Replenishment of naive T cells (thymus input)
- $N_0$ = 10^6 naive cells
- $V_0$ = 5x10^5 Tvm cells
- $k_{female}$ = 0.03
- $k_{male}$ = 0.03 × (1.1 / 0.7) = 0.047
- $s_{female}$ = 20,000 ($t$≤4); $s_{male}$×0.7 (4<$t$≤8); $s_{male}$×0.4 (8<$t$≤12); $s_{male}$×0.15 ($t$>12)
- $s_{male}$ = 5,000 ($t$≤4); $s_{male}$×0.3 (4<$t$≤8); $s_{male}$×0.15 (8<$t$≤12); $s_{male}$×0 ($t$>12)

$N_{t+1} = N_t \times (1-k)$
$C_t = 1-(1-k)^t$
Conversion (%) $= 100 \times (1-(1-k)^t)$
$k_{female} = 0.03$
$C_{female}(t) = 1-(1-0.03)^t$
$k_{male} = 0.03 \times (1.1 / 0.7) = 0.047$
$C_{male}(t) = 1 - (1 - 0.047)^t$

$N_{t+1} = N_t \times (1-k)$
$V_t+1 = V_t + k_{sex} \times N_t$
naive pool: $N_t = N_0 \times (1-k)^t$
Tvm pool: $V_t = V_0 + N_0 \times (1-(1-k)^t)$
$N_t + V_t = N_0 + V_0$

$N_{sex}(t+1) = N_{sex}(t) + s_{sex}(t) - k_{sex} \times N_{sex}(t)$
$V_{sex}(t+1) = V_{sex}(t) + k_{sex} \times N_{sex}(t)$

**Supplementary Data Fig. 1. Conceptual framework for cumulative naive CD8 T cell to virtual memory T cell conversion. a,** Definition of parameters. **b, *Cumulative conversion over time.*** To assess whether the observed sex differences in conversion propensity are sufficient to account for progressive divergence in naive CD8 T cell maintenance, we implemented a minimal discrete-time model of cumulative naive-to-Tvm conversion based on models of compounding. In this framework, a fixed fraction $k$ of the remaining naive pool converts to Tvm cells per time interval, such that the naive fraction evolves according to $N_{t+1} = N_t(1 - k)$, yielding $N_t = (1 - k)^t$. The cumulative fraction of converted cells is therefore $C_t = 1 - (1 - k)^t$. Based on published adoptive transfer experiments (White et al., *Nature Communications* 2016), we set $k_{female}$ = 0.03 per 3-week interval for females. To account for the increased frequency of conversion-prone cells in males, we scaled this rate by the observed ratio of CD5^high naive CD8 T cells **(Fig. 3f)**, yielding $k_{male}$ 0.03*(1.1/0.7) = 0.047. This minimal model demonstrates that modest differences in per-interval conversion rates produce substantial divergence in cumulative conversion over time. **c*, Naive reduction and Tvm accumulation (conversion driven only).*** We modeled naive CD8 T cell decline as a discrete-time process in which a fixed fraction $k$ of the naive pool converts to Tvm cells per interval, in the absence of thymic input. This yields $N_{t+1} = N_t(1 - k)$ and $V_{t+1} = V_t + k*N_t$, resulting in exponential decay of the naive pool and corresponding cumulative accumulation of Tvm cells. **d*, Naive reduction and Tvm accumulation (conversion + Thymic input).*** To incorporate sex-biased thymic support, we extended the conversion model by adding a time-dependent thymic input term to the naive compartment. Naive CD8 T cell dynamics were modeled as $N_{t+1} = N_t + s_t - k*N_t$, with corresponding Tvm accumulation given by $V_{t+1} = V_t + k*N_t$. Thymic input was modeled as a gradually declining function $s$ with more sustained thymic input in females compared to males.

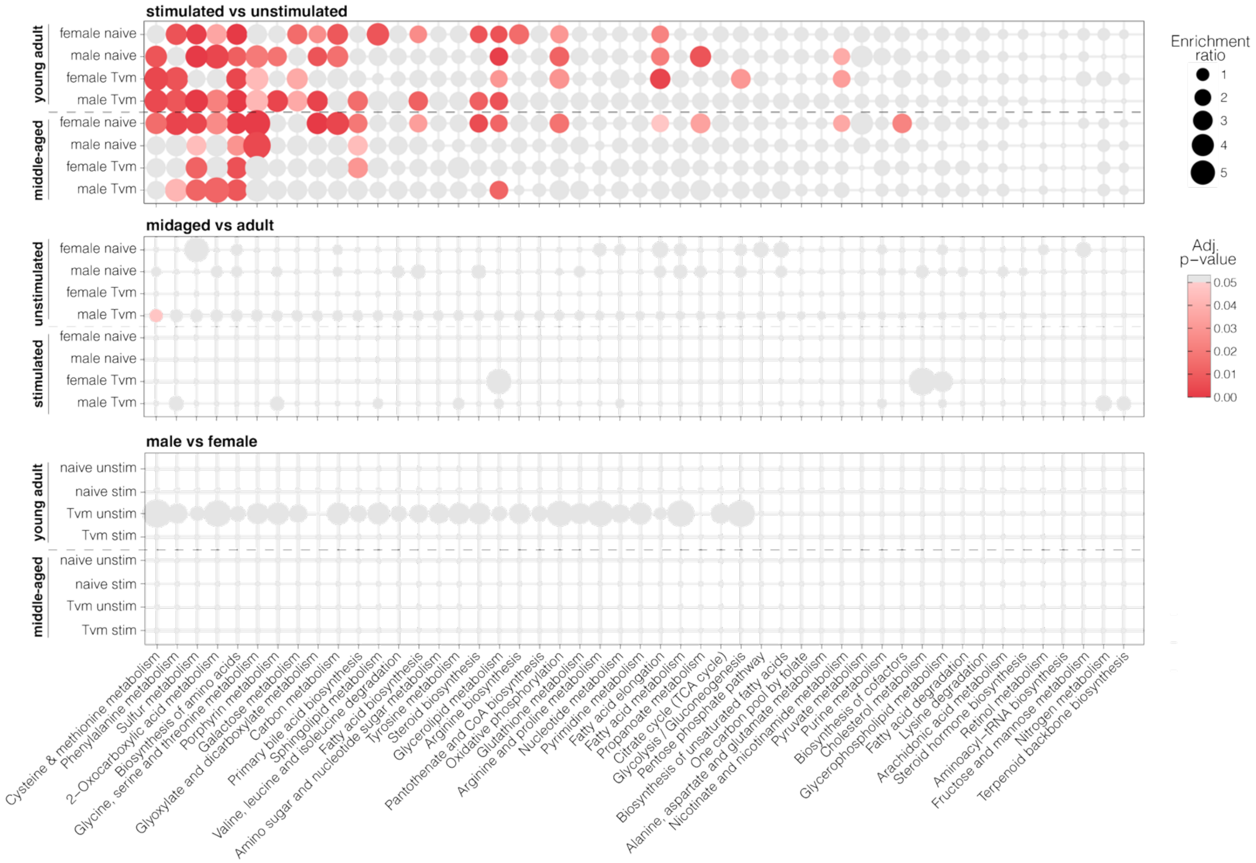


**Supplementary Data Fig. 2. KEGG metabolic pathway enrichment analysis.** Tvm and naive CD8 T cells from young adult and middle-age female and male mice were analyzed after 5 hours of CD3/CD28 in vitro stimulation. Dot size indicates enrichment ratio: (differentially expressed genes in pathway / total differentially expressed genes genes) / (all genes in pathway / all genes in genome); color indicates FDR-adjusted p-value (grey = not significant, red = adj. $p < 0.05$). Statistics calculated with BH-corrected hypergeometric test p-value.
